# Predator stimulus and habitat structure jointly shape antipredator behavior in frog species

**DOI:** 10.64898/2026.05.26.728007

**Authors:** Diogo B. Provete, Jessyca M. Citadini, Fernando R. Gomes

## Abstract

When encountering a predator, prey must choose between immobility and flight, a decision shaped by predator proximity, habitat structure, body size, and evolutionary history. Despite extensive work on optimal escape theory, few studies have jointly modelled both the decision to flee and the intensity of flight in a phylogenetic comparative framework. Here, we used a Bayesian phylogenetic hurdle log-normal model to analyze the probability of immobility and the conditional jump distance in 534 trials from 89 males across 17 Neotropical frog species (seven families), exposed to a simulated snake predator in three arenas of varying structural complexity. Physical contact with the predator (touch) was the strongest predictor, reducing immobility probability from 94% to 15% and increasing jump distance by 26%. Habitat complexity increased immobility (bush > leaf litter > empty), and frogs in open arenas jumped 31% farther. Larger-bodied species were substantially more likely to remain immobile but did not jump farther, indicating that body size determines strategy choice rather than locomotor magnitude. Phylogenetic signal was strong for both components (Pagel’s λ = 0.80 for jump distance; λ = 0.71 for immobility), with phylogeny accounting for 69–75% of variance. Species’ random effects formed a phylomorphospace in which fossorial species showed highest immobility and arboreal/torrent species favored active escape. Substantial individual-level variation in immobility tendency (25% of variance) provides a heritable substrate for ongoing selection. Frog antipredator strategies are jointly shaped by ecological context and evolutionary history; hurdle models offer a powerful framework for decomposing behavioral decisions.

**LAY SUMMARY:** When a predator approaches, prey must choose between staying still or fleeing — a decision depending on body size, habitat, and shared evolutionary history. We exposed 17 frog species to a snake in arenas of varying vegetation cover. Frogs fled when touched but mostly stayed still when only approached; larger species and those in dense vegetation were most likely to remain immobile. These antipredator strategies were strongly shaped by frogs’ shared ancestry.

## INTRODUCTION

Predation is not a single event, but a sequence of decisions distributed across multiple, partially independent stages: encounter, detection, identification, approach, subjugation, and consumption, each of which presents prey with distinct opportunities to interrupt the chain of events that ends with capture (Endler 1986; Lima and Dill 1990; Kikuchi et al. 2023). At each stage, different defenses become effective: pre-detection stages favor camouflage, background matching, and reduced movement, whereas post-detection stages reward escape performance, evasive trajectories, and, ultimately, defenses that mitigate harm during physical contact (Suraci et al. 2022; Whitford et al. 2019; Linke et al. 2025). Recent comparative analyses suggest that prey rarely succeed by relying on a single line of defense. Instead, defense portfolios distributed across the sequence appear to be the rule, rather than the exception (Kikuchi et al. 2023). The result is that predation success in many systems is remarkably low — often only 1–5% of attacks end in a kill (Whitford et al. 2019), because at every stage the prey can deploy a context-appropriate response that may break the sequence.

Within this framework, behavior occupies a privileged position: it operates as a filter between whole-organism performance capacity and the actual fitness consequences of an encounter (Garland and Losos 1994). An animal carrying a body plan capable of high locomotor performance does not necessarily express that performance during every predation event; the behavioral decision of whether, when, and how intensely to deploy that performance fundamentally shapes the relationship between morphology and survival. Consequently, the evolutionary pressures imposed by predation act not only on performance-relevant morphology, but also, and arguably more directly, on the behavioral rules that govern its deployment (Arnold 1983; Dickinson et al. 2000; Langerhans 2006).

Optimal escape theory (OET) formalizes these behavioral decisions by treating them as economic problems: prey should escape at the point where the cost of fleeing — in lost opportunities for foraging, mating, or thermoregulation, plus the energetic expenditure of locomotion itself — just balances the residual risk of remaining in place (Ydenberg and Dill 1986; Cooper and Frederick 2007; Cooper et al. 2015; Samia et al. 2016). A central insight of this theory (Cooper and Blumstein 2015) is that the optimal solution is rarely a single fixed behavior. Instead, it depends on a set of state variables: distance to the predator, distance to refuge, body condition, prior detection, which prey integrate to choose between two qualitatively different defensive modes. Cryptic strategies (immobility coupled with background-matching coloration, postures, or background-matching texture) minimize the probability of being detected and identified as prey but become useless once detection has occurred (Cooper et al. 2008). Active strategies (fleeing, deimatic displays, defensive secretions, or aggression) impose direct costs but maximize the probability of survival once cover is broken (Kikuchi et al. 2023). The transition between these modes is the central problem in animal antipredator decision-making.

Empirical studies across vertebrates broadly support the OET framework (Stankowich and Blumstein 2005). Lizards, the best-studied group (Samia et al. 2016), modulate flight initiation distance with predator approach speed, body size, refuge distance, and habitat structure. In fish and amphibians (Fleming and Bateman 2015), escape decisions are similarly sensitive to multiple risk and cost components, with the additional complication that ectotherm performance is strongly temperature-dependent. Across taxa, two factors emerge as particularly powerful predictors of strategy choice: the proximity and nature of the predator stimulus (which determines the stage of the predation sequence the prey perceives itself to be in; Kikuchi et al. 2023; Hemmi & Pfeil 2010), and the structural complexity of the immediate environment (which determines the relative payoffs of crypsis versus active escape; Merilaita et al. 1999; Wishingrad et al. 2014). Yet a complete empirical test requires manipulating both factors orthogonally and quantifying both components of the response: the decision to flee at all, and the intensity of escape once chosen in a phylogenetically informative sample.

Anurans (frogs and toads) provide an unusually tractable system for such a test. Their body plan, established at least by the Early Jurassic (Shubin and Jenkins 1995), is one of the most morphologically distinctive among tetrapods, with an elongated pelvis, fused postsacral vertebrae, and proportionally enormous hindlimbs that together specialize the skeleton for saltatorial locomotion (Emerson 1985; Reilly and Jorgensen 2011; Jorgensen and Reilly 2013; Buttimer et al. 2020). This morphological canalization is paired with a dramatic ecological radiation: contemporary anurans inhabit microhabitats ranging from open leaf litter and grasslands to fossorial chambers, riparian rocks, pond margins, and the canopies of tropical forests (Haddad et al. 2008; Buttimer et al. 2020). The interaction between body-plan conservation and microhabitat radiation has produced predictable patterns of phenotypic divergence: arboreal, torrent, and semi-aquatic species have evolved relatively longer hindlimbs and higher jumping performance than terrestrial and fossorial species (Gomes et al. 2009; Moen et al. 2013; Citadini et al. 2018; Leavey et al. 2023), with corresponding shifts in jumping power that scale negatively with body mass and reflect microhabitat-specific biomechanical trade-offs (Emerson 1978; Mendoza et al. 2020). Within arboreal species alone, hindlimb form–function relationships diverge across a span of ∼120 million years of evolution (Simon et al. 2025), and interspecific variation in take-off and landing kinematics reflects functional differentiation of the iliosacral articulation across pelvic morphotypes (Reilly et al. 2016a, b). Detailed kinematic studies further reveal how landing strategies and adhesive morphology have been fine-tuned to perch geometry (Abdala et al. 2022).

This rich understanding of how microhabitat has shaped morphology and performance in anurans contrasts sharply with the relative paucity of data on how it has shaped the behavioral rules that govern when and how that performance is deployed against predators. If behavior is genuinely a filter on the action of selection on performance (Arnold 1983; Garland and Losos 1994), then we should expect anuran antipredator strategies to have co-diversified with locomotor performance across microhabitat axes, with arboreal and torrent species relying more heavily on athletically demanding active escape (Figure 1), and terrestrial and fossorial species relying more on cryptic immobility, body posture, and chemical defenses (Williams et al. 2000; Toledo et al. 2011). Although a few field studies have shown that sympatric frog species differ markedly in the balance between freezing and fleeing (Cloyed and Eason 2015; Matich and Schalk 2019), and that body-size × microhabitat interactions shape individual escape decisions (Bateman and Fleming 2014), no study has tested these patterns jointly across a phylogenetically broad sample under standardized conditions.

**Figure 1.**
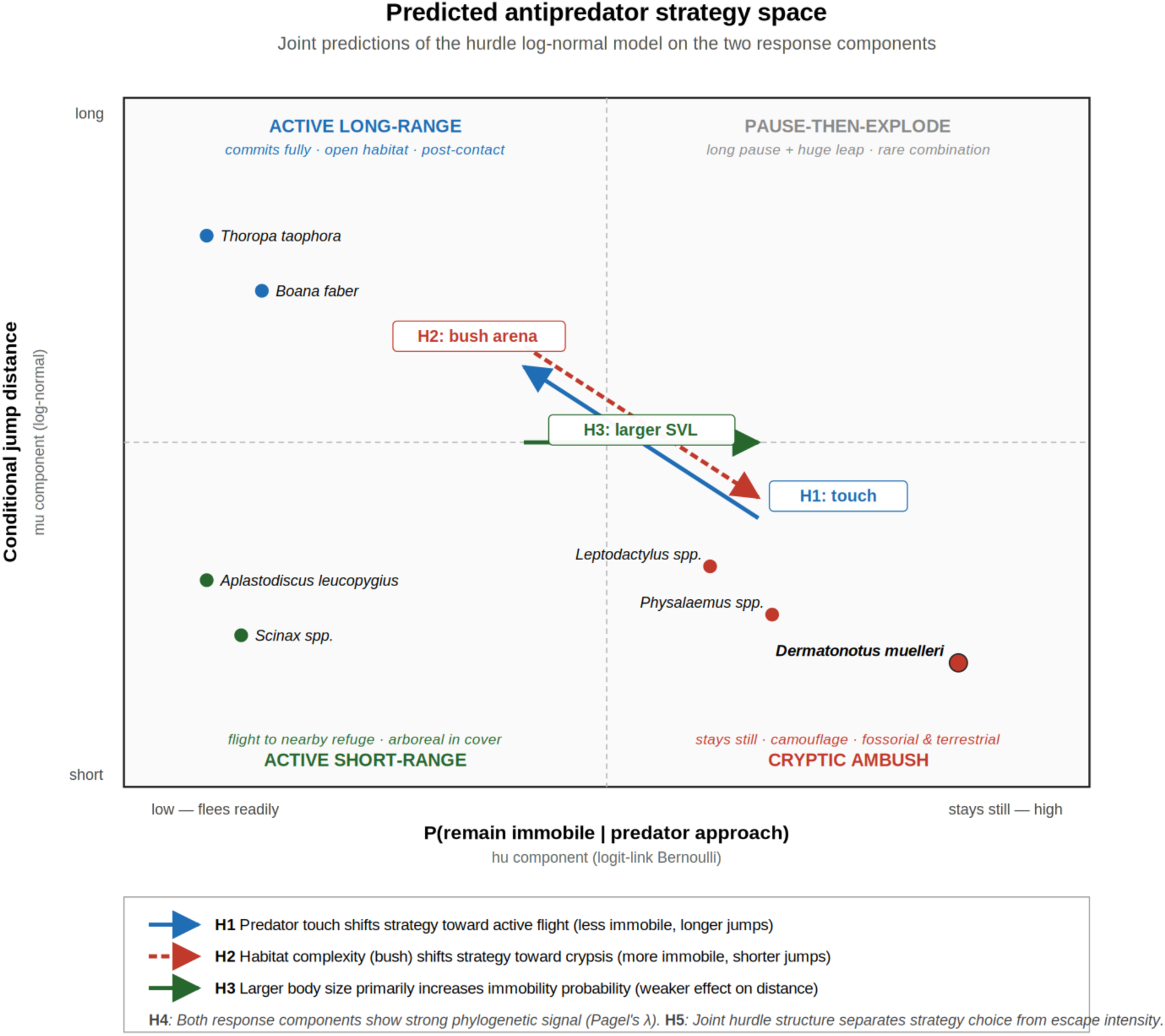
Predicted antipredator strategy space. Schematic hypothesis diagram illustrating the joint predictions of the hurdle log-normal model on the two response components. The x-axis represents conditional jump distance (*mu* component) and the y-axis represents immobility probability (*hu* component). The five hypotheses (H1–H5) are mapped onto predicted directions of effects. Exemplar species are positioned to illustrate expected ecological associations with microhabitat type.

Frogs deploy a remarkable diversity of antipredator behaviors. Toledo et al. (2011) catalogued at least 30 distinct defensive tactics, including immobility (often coupled with cryptic coloration) and active flight by jumping. A global database (Ferreira et al. 2019) extended this inventory to 2,953 records from 650 post-metamorphic species, classifying 12 antipredator mechanisms with 28 behavioral variations into three functional phases of the predation sequence: avoiding detection, preventing attack, and counterattacking after physical contact. Their analysis revealed that camouflage, immobility, and escape are widespread ancestral components of the anuran defense repertoire, whereas traits such as distress calls and interrupt calling are convergent across families (Ferreira et al. 2019). Empirical work on individual species has consistently shown that the choice among tactics is modulated by the predator stimulus and the habitat context. Tree frogs (*Ololygon hiemalis*) jumped shorter distances when exposed to a snake in their structurally complex natural microhabitat than in the open floor of a laboratory arena (Gomes et al. 2002), suggesting that anurans modulate locomotor effort according to refuge availability. Iberian green frogs (*Pelophylax perezi*) showed the same response to aquatic vegetation (Martín et al. 2005). At a finer behavioral scale, Japanese frogs switch from immobility to active fleeing as the approaching predator crosses a threshold distance, revealing that the strategy decision itself is dynamic and stimulus-driven (Nishiumi and Mori 2015). Túngara frogs (*Engystomops pustulosus*) even tailor their escape trajectory to the attack vector of the predator, fleeing away from terrestrial snakes, but moving toward bat models to undercut the aerial attack path (Bulbert et al. 2015). Experimental comparison of cryptic and aposematic frogs further showed that cryptic species rely heavily on immobility under simulated predator approach, while aposematic species are more likely to move, a behavioral component of warning signal display (Blanchette et al. 2017).

Snakes are among the main predators of frogs in the Neotropics, and many of the species we examine here have documented snake predators in the Atlantic Forest and adjacent biomes. Anurophagous colubrids and dipsadids, such as *Chironius* spp., *Erythrolamprus miliaris*, *Leptophis ahaetulla*, and *Thamnodynastes* spp. consume both terrestrial and arboreal frogs, including species in our sample (Marques et al. 2001; Eisfeld et al. 2021; Muscat et al. 2017; Oliveira et al. 2022). The predominance of snake predation makes the slow-approach + sudden-strike attack pattern a particularly relevant predator stimulus type for any comparative study of anuran antipredator decisions (Figure 1), and motivated our experimental design that exposed frogs to both visual approach and tactile contact by a snake stimulus.

Despite this body of work, three substantive gaps limit our understanding of how frog antipredator behavior has evolved. First, most empirical studies have focused on a single species or a small comparison of two or three sympatric taxa (e.g., Cloyed and Eason 2015; Bateman and Fleming 2014; Matich and Schalk 2019). Comparative inference about the evolution of strategy choice across the frog radiation requires a phylogenetically broader sample tested under standardized conditions, allowing among-species variation to be partitioned from environmental and design effects. Second, studies have typically analyzed one response variable at a time: either the binary decision of fleeing *versus* remaining still (often coded as a categorical outcome) or the magnitude of the escape jump conditional on flight, without modeling the joint structure of the two responses. Yet the two are biologically inseparable: they reflect, respectively, the decision of which branch of the predation sequence to engage, and the intensity of effort committed once that branch is chosen. A statistical framework that estimates them jointly, allowing covariates to act on each component independently, can reveal trade-offs and decision-logic patterns that single-component analyses cannot. Third, although the phylogenetic structure of frog morphology and locomotor performance is now well characterized (Citadini et al. 2018; Mendoza et al. 2020; Leavey et al. 2023; Simon et al. 2025), the phylogenetic structure of antipredator behavior remains largely unquantified. Recent methodological advances in multi-response phylogenetic mixed-effects models (Hadfield 2010; Bürkner 2017; Halliwell et al. 2024) make this estimation tractable, but they have not been applied to the joint estimation of behavioral strategy and locomotor intensity.

Here, we address these three gaps with a comparative experimental study of antipredator behavior in 17 species of Neotropical frogs from seven families and spanning six microhabitats. We exposed individuals to a fully-crossed combination of three arena types varying in structural complexity (empty, leaf litter, and bush vegetation) and two predator stimuli representing successive stages of the predation sequence (visual approach by a snake stimulus, simulating pre-contact detection; and tactile touch by the same stimulus, simulating physical capture attempt). We then analyzed the resulting behavioral data with a Bayesian phylogenetic hurdle log-normal model that jointly estimates two response components: the probability of remaining immobile (a binary decision, modelled with a logit-link Bernoulli component), and the conditional jump distance given that flight occurred (modelled with a log-normal component). Both components incorporate phylogenetic random effects, with covariance structure derived from a recent species-level time-calibrated tree (Portik et al. 2023), as well as individual-level random effects to account for repeated trials.

The joint hurdle structure (Cragg 1971) allows us to test a set of hypotheses that bear on both the strategy-choice and the intensity components of escape behavior (Table 1; Figure 1). Specifically, we predicted that:

**Table 1.**
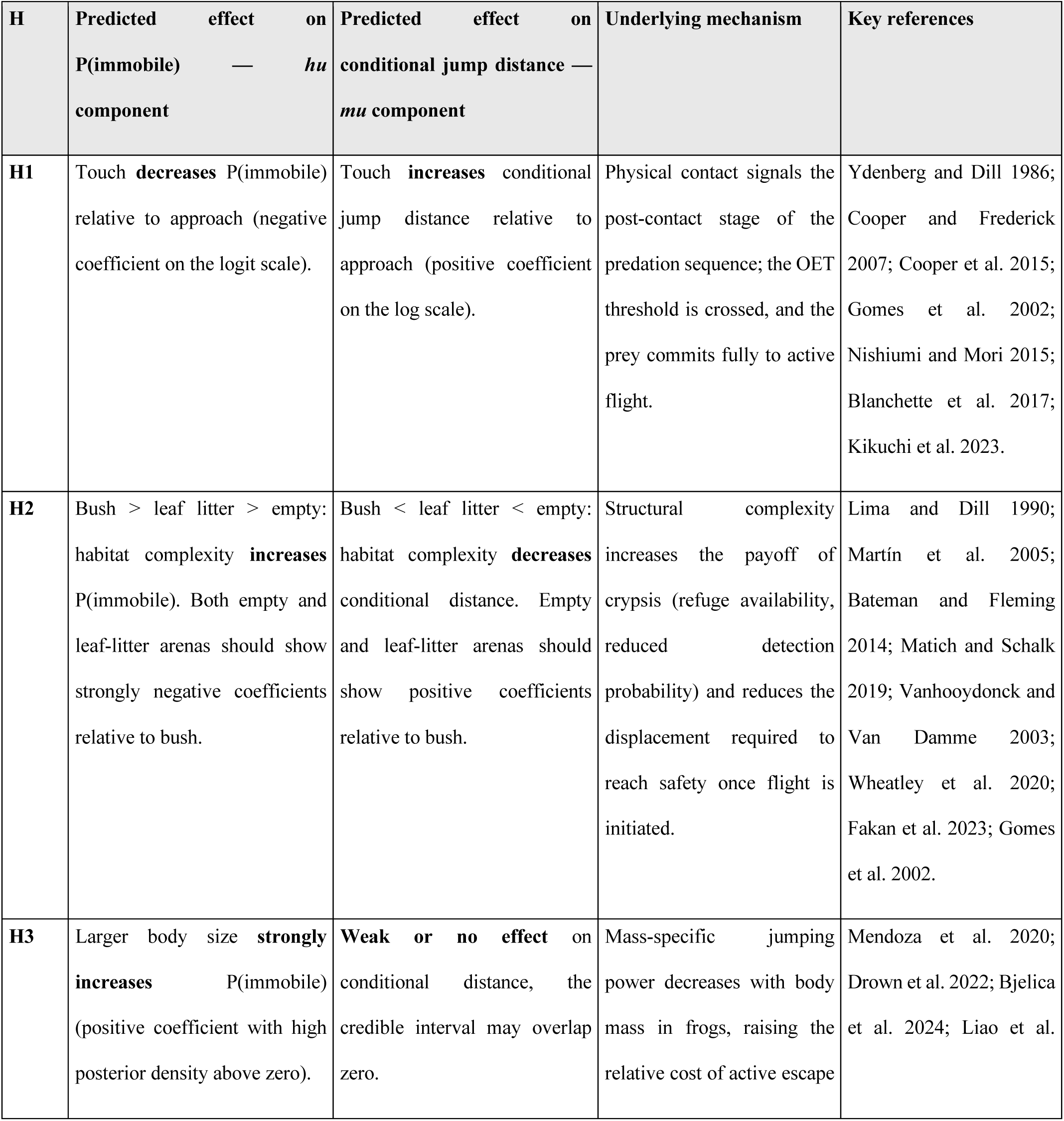
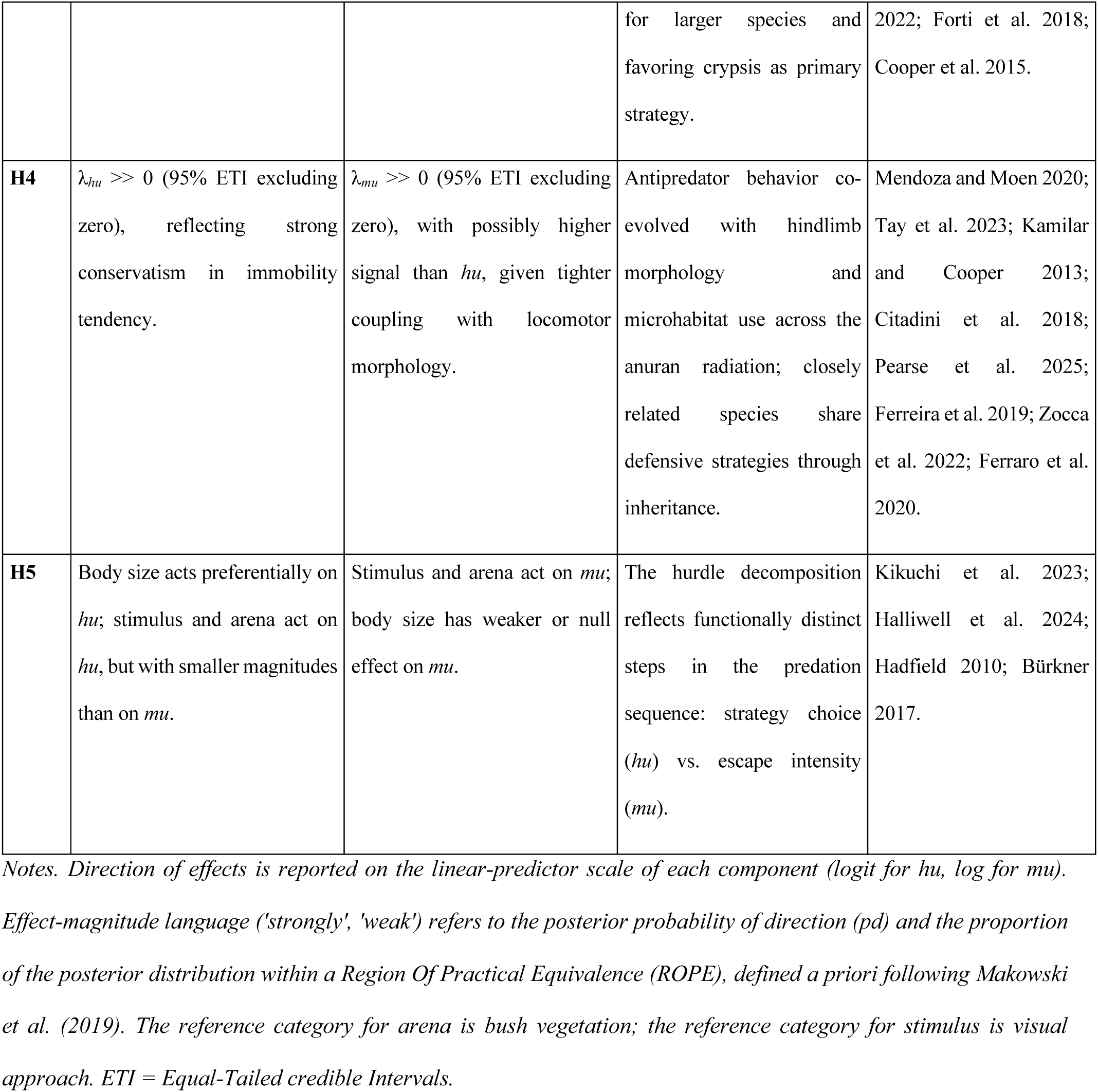
Hypotheses tested in this study, mapped onto the two response components of the joint Bayesian phylogenetic hurdle log-normal model: probability of remaining immobile under predator approach (*hu* component, logit-link Bernoulli) and conditional jump distance given that flight occurred (*mu* component, log-normal). Each hypothesis specifies the expected direction of effects, the underlying biological mechanism, and the theoretical or empirical basis.

| H | Predicted effect on P(immobile) — *hu* component | Predicted effect on conditional jump distance — *mu* component | Underlying mechanism | Key references |
| --- | --- | --- | --- | --- |
| H1 | Touch **decreases** P(immobile) relative to approach (negative coefficient on the logit scale). | Touch **increases** conditional jump distance relative to approach (positive coefficient on the log scale). | Physical contact signals the post-contact stage of the predation sequence; the OET threshold is crossed, and the prey commits fully to active flight. | Ydenberg and Dill 1986; Cooper and Frederick 2007; Cooper et al. 2015; Gomes et al. 2002; Nishiumi and Mori 2015; Blanchette et al. 2017; Kikuchi et al. 2023. |
| H2 | Bush > leaf litter > empty: habitat complexity **increases** P(immobile). Both empty and leaf-litter arenas should show strongly negative coefficients relative to bush. | Bush < leaf litter < empty: habitat complexity **decreases** conditional distance. Empty and leaf-litter arenas should show positive coefficients relative to bush. | Structural complexity increases the payoff of crypsis (refuge availability, reduced detection probability) and reduces the displacement required to reach safety once flight is initiated. | Lima and Dill 1990; Martín et al. 2005; Bateman and Fleming 2014; Matich and Schalk 2019; Vanhooydonck and Van Damme 2003; Wheatley et al. 2020; Fakan et al. 2023; Gomes et al. 2002. |
| H3 | Larger body size **strongly increases** P(immobile) (positive coefficient with high posterior density above zero). | **Weak or no effect** on conditional distance, the credible interval may overlap zero. | Mass-specific jumping power decreases with body mass in frogs, raising the relative cost of active escape | Mendoza et al. 2020; Drown et al. 2022; Bjelica et al. 2024; Liao et al. |
|  |  |  | for larger species and favoring crypsis as primary strategy. | 2022; Forti et al. 2018; Cooper et al. 2015. |
| **H4** | $\lambda_{hu} \gg 0$ (95% ETI excluding zero), reflecting strong conservatism in immobility tendency. | $\lambda_{mu} \gg 0$ (95% ETI excluding zero), with possibly higher signal than $hu$, given tighter coupling with locomotor morphology. | Antipredator behavior co-evolved with hindlimb morphology and microhabitat use across the anuran radiation; closely related species share defensive strategies through inheritance. | Mendoza and Moen 2020; Tay et al. 2023; Kamilar and Cooper 2013; Citadini et al. 2018; Pearse et al. 2025; Ferreira et al. 2019; Zocca et al. 2022; Ferraro et al. 2020. |
| **H5** | Body size acts preferentially on $hu$; stimulus and arena act on $hu$, but with smaller magnitudes than on $mu$. | Stimulus and arena act on $mu$; body size has weaker or null effect on $mu$. | The hurdle decomposition reflects functionally distinct steps in the predation sequence: strategy choice ($hu$) vs. escape intensity ($mu$). | Kikuchi et al. 2023; Halliwell et al. 2024; Hadfield 2010; Bürkner 2017. |

Notes. Direction of effects is reported on the linear-predictor scale of each component (logit for $hu$ , log for $mu$ ).
Effect-magnitude language ('strongly', 'weak') refers to the posterior probability of direction ( $pd$ ) and the proportion of the posterior distribution within a Region Of Practical Equivalence (ROPE), defined a priori following Makowski et al. (2019). The reference category for arena is bush vegetation; the reference category for stimulus is visual approach. ETI = Equal-Tailed credible Intervals.

### (H1) Predator stimulus drives strategy switching

Touch should reduce the probability of remaining immobile and should increase the conditional jump distance, both relative to visual approach, reflecting the transition from pre-contact crypsis to post-contact active escape predicted by OET and the predation-sequence framework.

### (H2) Habitat structural complexity reshapes the optimal strategy

Structurally complex arenas (bush > leaf litter > empty) should increase the probability of immobility and decrease the conditional jump distance, because complexity makes crypsis more effective and reduces the displacement required to reach refuge.

### (H3) Body size determines strategy choice

Larger species should show higher probabilities of immobility, reflecting both the higher relative cost of whole-body locomotion at large size and the established pattern that mass-specific jumping power decreases with body mass in anurans (Mendoza et al. 2020).

### (H4) Antipredator behavior is phylogenetically structured

Both response components should show substantial phylogenetic signal (Pagel’s λ >> 0), reflecting the co-diversification of behavior and morphology across the anuran microhabitat radiation (Buttimer et al. 2020). This prediction is consistent with phylogenetic signal estimates reported for anuran locomotor performance (K ≈ 0.7–0.8 for jumping velocity, energy, and power; Mendoza and Moen 2020) and with direct evidence of conservatism in antipredator behavior from clade-level comparative studies of horned frogs (Zocca et al. 2022) and leiuperine frogs (Ferraro et al. 2020).

### (H5) Strategy choice and escape intensity are partially independent

Joint estimation should reveal that some covariates (e.g., body size) act preferentially on the strategy-choice component, whereas others (e.g., stimulus type) act on both, a pattern obscured by single-response analyses, but expected if the two response dimensions reflect functionally distinct steps in the predation sequence.

By integrating a phylogenetically broad experimental sample with a joint Bayesian modelling framework, our study provides the first comparative test of how multiple factors influence both the strategic and the kinematic components of frog antipredator behavior, and the first quantitative estimate of phylogenetic signal in anuran escape decision-making. The results bear on three broader questions: 1) how prey deploy defenses across stages of the predation sequence (Kikuchi et al. 2023); 2) how morphological canalization and behavioral lability interact in evolutionarily diverse clades (Garland and Losos 1994; Simon et al. 2025); and 3) how the phylogenetic structure of behavior compares with that of the morphological and performance traits with which it co-evolves.

## MATERIALS AND METHODS

### Field work and sampling

Males from 17 frog species from seven families were collected by JMC in several localities in Southeastern Brazil (Figure S1) from November 2013 to December 2015 (Table S1). Each species was classified into six microhabitat types: aquatic (living predominantly in or near water), arboreal (living above the ground on vegetation), fossorial (living in burrows), terrestrial (living on the ground), cryptozoic (living predominantly amid leaf litter and burrowing habits), and torrent (living closely associated with rocky, fast-flowing streams) following Haddad et al. (2008).

### Animal maintenance in the laboratory

Specimens were placed in plastic boxes and transported to the laboratory by JMC, where they were housed individually in terraria: 13 × 30 × 28 cm for smaller species, and 27 × 45 × 32 cm for larger species. Each terrarium contained fragments of vegetation, stones, and freshwater permanently available; animals were fed weekly with small live cockroaches. All experiments were carried out two weeks after animals were captured. Terraria were maintained in a climate-controlled room at 25 °C (± 1 °C) and 12:12 h dark:light cycle.

We used *Erythrolamprus miliaris* (Serpentes: Colubridae: Dipsadinae), a medium-sized, semi-aquatic, diurnal–nocturnal snake widely distributed in the state of São Paulo (Uetz et al. 2026), to quantify the antipredator response. This species is a natural predator of frogs, usually associated with water bodies (Marques et al. 2001). All snakes (N = 8) were maintained in separate 27 × 45 × 32 cm transparent plastic boxes with fragments of vegetation, stones, resting places, and fresh water, fed weekly with small live fishes, in a climate-controlled room at 25 °C (± 1 °C) and 12:12 h dark:light cycle.

### Experimental design

Experiments were carried out by JMC supervised by FRG during the night (20:00–01:00 h), since most species are nocturnal (Haddad et al. 2008). The only exception was the diurnal *Hylodes asper*, for which experiments were conducted during the day (14:00–16:00 h). During experiments, the observer (JMC) used a headlamp with red light (500 lumens). Simulated predation trials were conducted in an arena of 200 × 90 × 70 cm in a climate-controlled room at 25 °C (± 1 °C). Each frog was tested in three arenas with distinct structural complexity: (1) completely empty; (2) leaf litter covering the floor; (3) leaf litter and plastic bushes 50 cm height, spaced 10 cm apart covering the whole floor (Figure S2). The order of tests in different arenas was randomized for each frog, with a resting day between experimental rounds to minimize handling effects.

Each frog was placed in the arena through a small lateral door, positioned 5 cm from the entrance (Figure S2). After 30 min of habituation, we carefully placed the snake near the frog or at the posterior end of the frog (Gomes et al. 2002). For the tests, snakes had the mouth sealed using tape to prevent predation, while allowing tongue-flicking. A rubber ring was fitted around the snake, which was then tethered to a fishing rod (100 cm) using 10 cm of string. Frogs exhibited several behavioral responses to the approach of the snake, including remaining immobile, performing antipredator postures, or jumping away from the snake. Behavioral responses were recorded as soon as the snake was released. The snake consistently moved its head toward the frogs and tongue flicked. Three consecutive contacts were allowed between frogs and the snake before measuring antipredator behavior. Then, the distance of the first jump was measured to the nearest 1 cm (Gomes et al. 2002).

Collection permits were issued by ICMBio/SISBIO (# 49250-1) and Instituto Florestal de São Paulo (# 611/2015 for Parque Estadual Intervales). All laboratory procedures were approved by the IACUC of Universidade de São Paulo (protocol CEUA # 192/2013). Snakes used as predator stimuli were borrowed from Instituto Butantan and returned after the study. Frogs were maintained in individual terraria for the minimum duration necessary and were returned to their collection sites after experiments. No animals were harmed during trials; snake mouths were sealed to prevent predation during staged encounters, following ASAB/ABS guidelines for studies involving predation (ASAB Ethical Committee/ABS Animal Care Committee 2023). We adhered to the ARRIVE 2.0 guidelines for reporting animal research.

### Phylogenetic tree

We pruned the time-calibrated frog phylogeny of Portik et al. (2023) that includes 5,242 species, inferred from hundreds of genetic loci by combining phylogenomic data with a novel supermatrix, to retain only the 17 focal species. Nomenclature was updated to current synonymies following Frost (2024). The resulting 17-species ultrametric tree with branch lengths in divergence time (My) is deposited at Zenodo (Anonymous 2026).

### Statistical analysis

Statistical methods are described following Davis and Kay (2023) and the Bayesian Analysis Reporting Guidelines (BARG) of Kruschke (2021). Posterior distributions are summarized by the posterior mean ± SD and 95% Equal-Tailed credible Intervals (ETI; quantile-based 2.5–97.5 percentiles).

#### Stage 1 — Observation model

We modelled the antipredator response of each individual trial as a two-component hurdle log-normal model (Cragg 1971; Bürkner 2017, 2018; Halliwell et al. 2024). This formulation was motivated by the observation that 41% of all 534 trials resulted in immobility (jump distance = 0 cm), representing structural zeros: any individual that remained immobile necessarily produced a zero regardless of arena or stimulus conditions (Figure S10 and S11). The hurdle component (*hu*) models the probability of immobility (P[Y = 0]), while the log-normal component (*mu*) models the conditional distribution of jump distance given that the individual jumped (Y > 0). The *mu* component uses an identity link on the log scale; the *hu* component uses a logit link (Bürkner, 2018). Posterior predictive checks confirmed adequate recovery of both the proportion of structural zeros and the conditional density of non-zero distances (Figure S5).

Within-species adjusted repeatability (R) for each response component was estimated from the posterior variance componentes, following Nakagawa and Schielzeth (2010). For the log-normal component, R_mu_ = σ²_ind_/(σ²_ind_ + σ²_res_). For the Bernoulli hurdle component, which lacks an observation-level residual, we used the logistic-scale additive overdispersion variance π²/3 ≈ 3.29 as the denominator term (Nakagawa and Schielzeth 2010; Stoffel et al. 2017): R_hu_ = σ²_ind_/(σ²_ind_ + π²/3). Posterior distributions of R were obtained by computing R at each MCMC draw, fully propagating uncertainty from the variance components.

#### Stage 2 — Linear predictors

Both components shared the same covariate structure. Fixed effects included the full factorial interaction between arena type (three levels: bush [reference], empty, leaf litter) and predator stimulus (two levels: approach [reference], touch), and log-transformed grand-mean-centered snout–vent length (log SVLc) as an individual-level body-size covariate (see Anonymous 2026).

#### Stage 3 — Random-effects structure

Two nested random intercepts were specified for each component: (i) a phylogenetic random intercept for species, structured by the phylogenetic variance–covariance matrix **A** computed from the pruned tree, normalizing to a correlation matrix makes the phylogenetic SD directly comparable to other variance components (Housworth et al. 2004; Hadfield and Nakagawa 2010); and (ii) an individual random intercept for the repeated-measures design (89 frogs, each tested across three arenas). Species random effects follow multivariate normals structured by the phylogenetic covariance; individual intercepts are independent normals (see Anonymous 2026).

#### Stage 4 — Prior distributions and prior predictive check

Weakly informative priors were chosen, following Gelman et al. (2008) and Bürkner (2017). For the *mu* component: intercept ∼ N(5, 1.5²) [exp(5) ≈ 148 cm; spans biologically realistic jump distances]; fixed-effect slopes ∼ N(0, 1²); residual SD ∼ half-*t*(3, 0, 0.5); phylogenetic and individual SDs ∼ half-*t*(3, 0, 1). For the *hu* component: intercept ∼ N(0, 1.5²); fixed-effect slopes ∼ N(0, 2.5²) [standard weakly informative prior for logistic regression on standardized predictors; Gelman et al. 2008]; SDs ∼ half-*t*(3, 0, 1). Prior predictive checks confirmed that the prior generates plausible zero proportions (10–90%) and log-scale distances consistent with the observed range (see Anonymous 2026).

##### Computation and convergence (BARG steps 2–3)

Models were fitted with `brms` v. 2.22 (Bürkner 2017, 2018) using `Stan` (Carpenter et al. 2017) via `CmdStanR` (Gabry and Česnovar 2023). We ran four parallel MCMC chains with 6,000 iterations each (warm-up = 2,000; thinning = 2), yielding 8,000 post-warm-up draws. All R^ ≤ 1.001 (threshold: 1.01; Vehtari et al. 2021) and all Bulk-ESS and Tail-ESS > 2,000 (see Figures S3 and S4).

##### Prior sensitivity analysis (BARG step 5)

We conducted a prior sensitivity analysis using the `priorsense` R package (Kallioinen et al. 2024), which implements power-scaling of the prior and likelihood. Posteriors were insensitive to ± 25% prior perturbations for all *mu* parameters. Likelihood-scaling flags for some *hu* parameters reflect strong data informativeness — a desirable property confirming that the likelihood, not the prior, drives inference for these effects (see Figures S5 and S6).

##### Phylogenetic signal

Pagel’s λ (Pagel 1999) for each component was derived from the posterior variance components: λ*_mu_* = σ²_phylo_ / (σ²_phylo_ + σ²_ind_ + σ²), and analogously for λ*_hu_* (without σ², since the hurdle component is binomial). λ = 1 corresponds to Brownian motion; λ = 0 to phylogenetic independence (Pagel 1999; Pearse et al. 2025).

##### Model comparison

We compared M1 (hurdle log-normal, N = 534) against M2 (log-normal restricted to non-zero observations, N = 314) using leave-one-out cross-validation (LOO-CV; Vehtari et al. 2017). Pareto-*k* diagnostics confirmed reliable LOO estimates for both models (all *k* < 0.7).

##### Phylogenetic visualization

To visualize the distribution of behavioral strategies across the phylogeny, we constructed a behavioral phylomorphospace (Adams & Collyer 2019), projecting the phylogeny into a two-dimensional space defined by the posterior median species random effects: the *mu* RE (x-axis: deviation in log-distance from the population mean) and the *hu* RE (y-axis: deviation in logit-immobility). Internal node positions were estimated assuming a Brownian motion model of evolution in `phytools` (Revell 2024). Posterior 95% ETIs are shown as orthogonal error bars.

##### Software

All analyses were conducted in `R` 4.5 (R Core Team 2025) using `brms` (Bürkner 2017, 2018), `tidybayes` (Kay 2024), `bayesplot` (Gabry et al. 2019), `posterior` (Bürkner et al. 2022), `priorsense` (Kallioinen et al. 2024), `loo` (Vehtari et al. 2017), `ape` (Paradis and Schliep 2019), `phytools` (Revell 2024), `ggtree` (Yu et al. 2017), and the `tidyverse` (Wickham et al. 2019). A fully reproducible Quarto document with all code, data, model objects, and supplementary diagnostics is available in Anonymous (2026).

## RESULTS

We analyzed 534 individual trials from 89 males of 17 frog species from seven families (Table S1). Each frog was exposed to six treatment combinations (3 arenas × 2 stimuli), generating six observations per individual. Of 534 trials, 220 (41.2%) resulted in immobility (jump distance = 0 cm; structural zeros), and 314 (58.8%) resulted in active escape. The approach stimulus produced 114 of 267 jumping trials (42.7%), while touch produced 200 of 267 jumping trials (74.9%), confirming the strong stimulus effect already visible in the raw data.

### Immobility probability (hu component)

The reference condition (bush arena, approach stimulus, grand-mean SVL) was characterized by a posterior probability of immobility of 94.3% (95% ETI: [73.3, 99.2%]). All estimates below correspond to population-level predictions.

#### Predator stimulus (H1)

Physical contact with the predator (touch *vs.* approach) was the strongest effect in the model, reducing the log-odds of immobility by 4.55 ± 0.53 units (95% ETI: [−5.65, −3.56]; pd > 99.9%, ROPE = 0.0%). In the bush arena, this corresponds to a drop from 94.3% immobility under approach to 14.6% under touch, a reduction of ∼ 79% (Figure 2). This result confirms Hypothesis 1: animals responded with active escape to physical contact and with immobility to visual approach.

**Figure 2.**
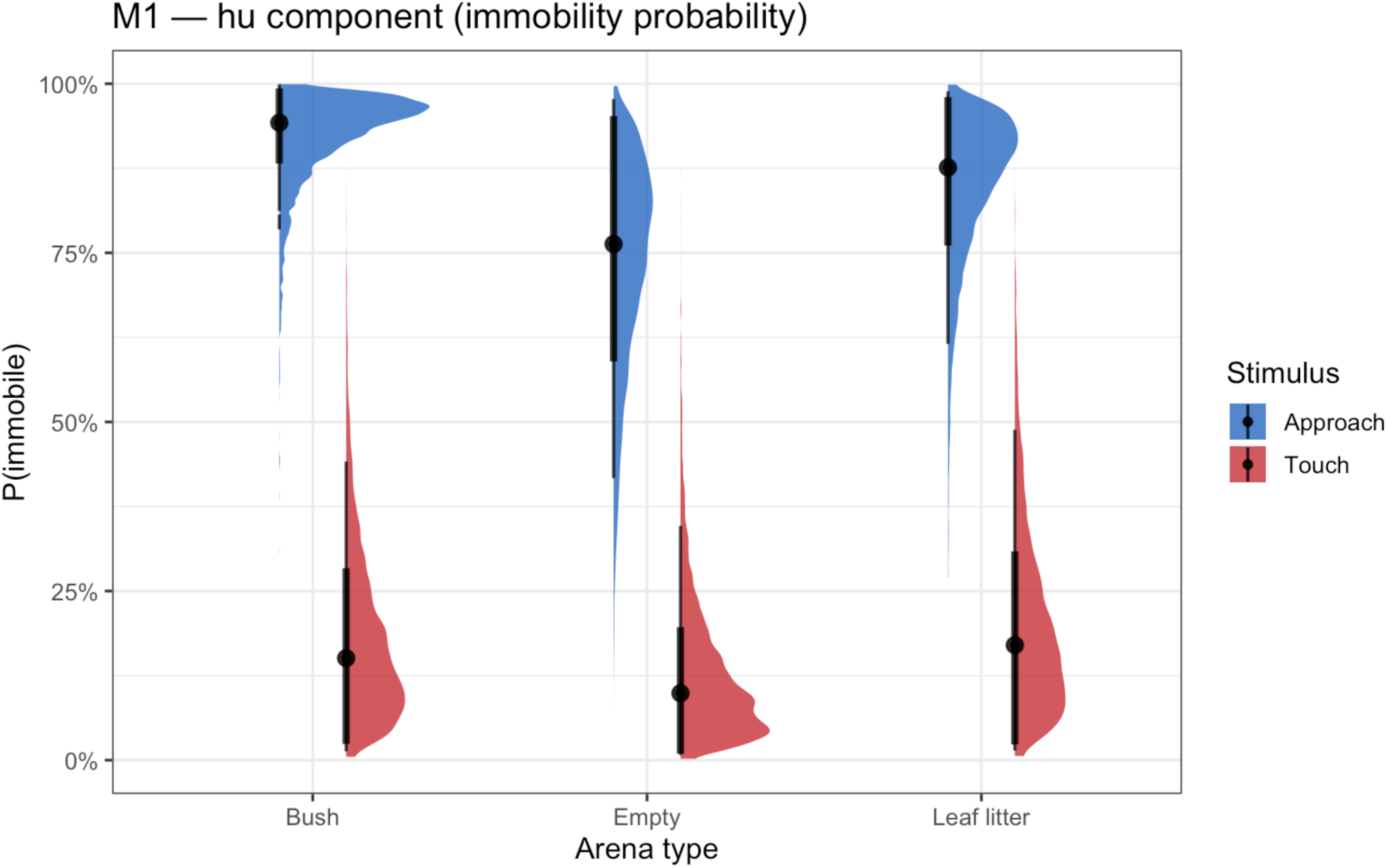
Marginal posterior predictions for the probability of remaining immobile (*hu* component) across arena types and predator stimuli. Violin and half-eye plots show the posterior predictive distribution for each treatment combination. Points represent individual posterior draws; thick bars indicate 66% and 95% credible intervals. Note the dramatic reduction in immobility from ∼94% under approach to ∼15% under touch in the bush arena.

#### Arena type (H2)

Compared with the bush arena, open environments reduced immobility. In the empty arena: β = −1.62 ± 0.39 (95% ETI: [−2.40, −0.86]; pd > 99.9%, ROPE = 0.0%), corresponding to a predicted P(immobile | approach) of 77.2%. In leaf litter: β = −0.84 ± 0.39 (95% ETI: [−1.62, −0.08]; pd = 98.3%, ROPE = 0.0%), corresponding to 87.6%. Both effects are clearly directional and outside the ROPE, partially confirming Hypothesis 2: habitat complexity increases immobility probability.

#### Arena × stimulus interaction

The interaction terms were positive in both cases: empty arena × touch: β = +1.15 ± 0.63 (95% ETI: [−0.09, +2.39]; pd = 96.7%); leaf litter × touch: β = +0.99 ± 0.62 (95% ETI: [−0.23, +2.20]; pd = 94.6%). Although credible intervals include zero, the consistently positive direction (pd > 94%) suggests a tendency for the touch effect to be partially attenuated in more open arenas, possibly because animals already had higher escape propensity in open environments.

#### Body size (H3)

Larger species had substantially higher probabilities of immobility (β = +2.37 ± 0.71; 95% ETI: [+1.00, +3.76]; pd > 99.9%, ROPE = 0.0%). A one-SD increase in body size (log scale) shifted the log-odds of immobility by +2.37 units, corresponding to a change from ∼77% to ∼96% immobility at the population-level intercept. This result indicates that larger-bodied species rely more heavily on crypsis as a primary antipredator strategy.

### Conditional jump distance (mu component)

The 314 non-zero jump distances (N = 89 individuals across 3 arenas × 2 stimuli) ranged from 3 to 1,009 cm (median: 252 cm). The reference condition posterior intercept corresponded to a predicted median conditional distance of 168 cm (95% ETI: [77, 354 cm]).

#### Predator stimulus (H1)

Touch produced 25.7% longer conditional jumps than approach (β = +0.229 ± 0.086; 95% ETI: [+0.060, +0.396]; pd = 99.5%, ROPE = 0.0%; predicted median: 211 cm). This directional shift, opposite to the immobility result, confirms that individuals who escape actively under touch commit more forcefully to flight (Figure 3).

**Figure 3.**
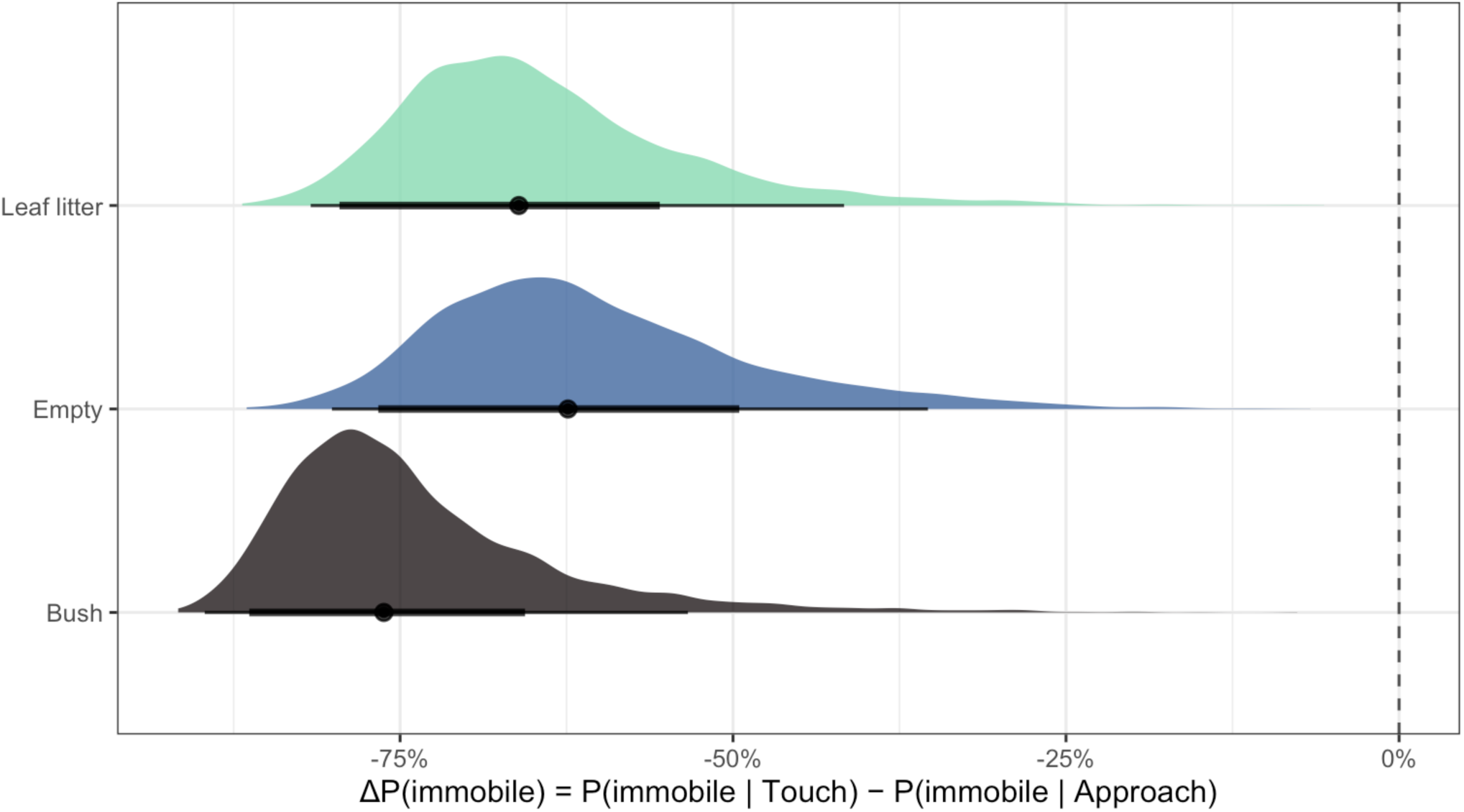
Posterior contrast densities for the touch effect on immobility probability across arena types. Each density curve represents the posterior distribution of the difference in P(immobile) between approach and touch conditions within a given arena. Negative values indicate that touch reduces immobility. All arenas show unambiguously negative contrasts (pd > 99.9%), confirming that physical contact triggers the switch to active escape regardless of habitat complexity.

#### Arena type (H2)

Conditional jump distances were ∼31% longer in the empty arena (β = +0.272 ± 0.092; 95% ETI: [+0.092, +0.450]; pd = 99.8%, ROPE = 0.0%) and leaf litter (β = +0.270 ± 0.096; 95% ETI: [+0.083, +0.457]; pd = 99.7%, ROPE = 0.0%) relative to the bush. Despite also showing higher immobility probability in the bush, individuals that escaped in complex habitats actually jumped shorter distances, consistent with the hypothesis that structural complexity limits locomotor performance or shortens optimal escape distances (Hypothesis 2).

#### Arena × stimulus interaction

Interactions for conditional distance were near zero: empty × touch: β = +0.022 ± 0.107 (95% ETI: [−0.187, +0.234]; pd = 57.8%); leaf litter × touch: β = −0.155 ± 0.111 (95% ETI: [−0.371, +0.060]; pd = 91.8%). The arena effects on jump distance are therefore consistent regardless of stimulus type.

#### Body size (H3)

Log SVL showed a positive, but uncertain association with conditional jump distance (β = +0.246 ± 0.182; 95% ETI: [−0.121, +0.606]; pd = 91.3%). The 95% ETI crosses zero, but the pd of 91.3% and near-zero ROPE percentage indicate a consistent positive tendency: larger species tend to jump farther when they escape, but the effect is not well constrained. This contrasts with the unambiguously strong positive effect of body size on immobility probability, suggesting that size primarily determines strategy choice, rather than locomotor magnitude *per se* (H5).

### Variance components and phylogenetic signal (H4)

Variance partitioning revealed distinct structures for the two model components (Figure 5). For the *mu* component, fixed effects explained 12.3% of total variance, phylogeny 69.3%, individual 1.1%, and residual 17.3%. For the *hu* component, phylogeny accounted for 75.1% of variance and individual identity 24.9%.

Pagel’s λ was high for both components. For log-distance, λ*_mu_* ≈ 0.80 (95% ETI: [0.70, 0.94]), indicating that ∼80% of between-species variation in conditional jump distance is structured by phylogenetic relatedness (Figure S7). For immobility logit, λ*_hu_* ≈ 0.71 (95% ETI: [0.55, 0.90]), reflecting similarly strong phylogenetic signal in immobility strategy.

Species-level variance was substantially larger for the *hu* component than for the *mu* component (SD_species,hu_ = 1.813 ± 0.481 vs. SD_species,mu_ = 0.661 ± 0.130), confirming that species differ more markedly in immobility tendency than in conditional jump distance. Individual-level variance in immobility was also substantial (SD_ind,hu_ = 1.145 ± 0.274), indicating meaningful within-species individual variation in defensive strategy. Within-species adjusted repeatability was moderate for immobility tendency (R_hu_ = 0.30 [0.12, 0.48]), but low for conditional jump distance (R_mu_ = 0.06 [0.00, 0.18]), indicating that individual frogs are consistent in which strategy they deploy, but not in the intensity of active escape.

### Species-level behavioral strategies

The behavioral phylomorphospace (Figure 4a) placed species into distinct regions of a two-dimensional behavioral strategy space. *Dermatonotus muelleri* (fossorial; N = 5 individuals, 30 trials) had the most extreme *hu* random effect, with the 95% ETI entirely above zero, confirming it as substantially more immobile than the population mean. *Boana faber* (arboreal) and *Thoropa taophora* (torrent) occupied the active-escape quadrant. Arboreal species (*Boana* spp., *Scinax* spp., *Aplastodiscus leucopygius*) consistently fell below the immobility axis, confirming that arboreal species rely less on crypsis. The annotated phylogeny (Figure 4b) further shows how the two behavioral dimensions are distributed across clades.

**Figure 4.**
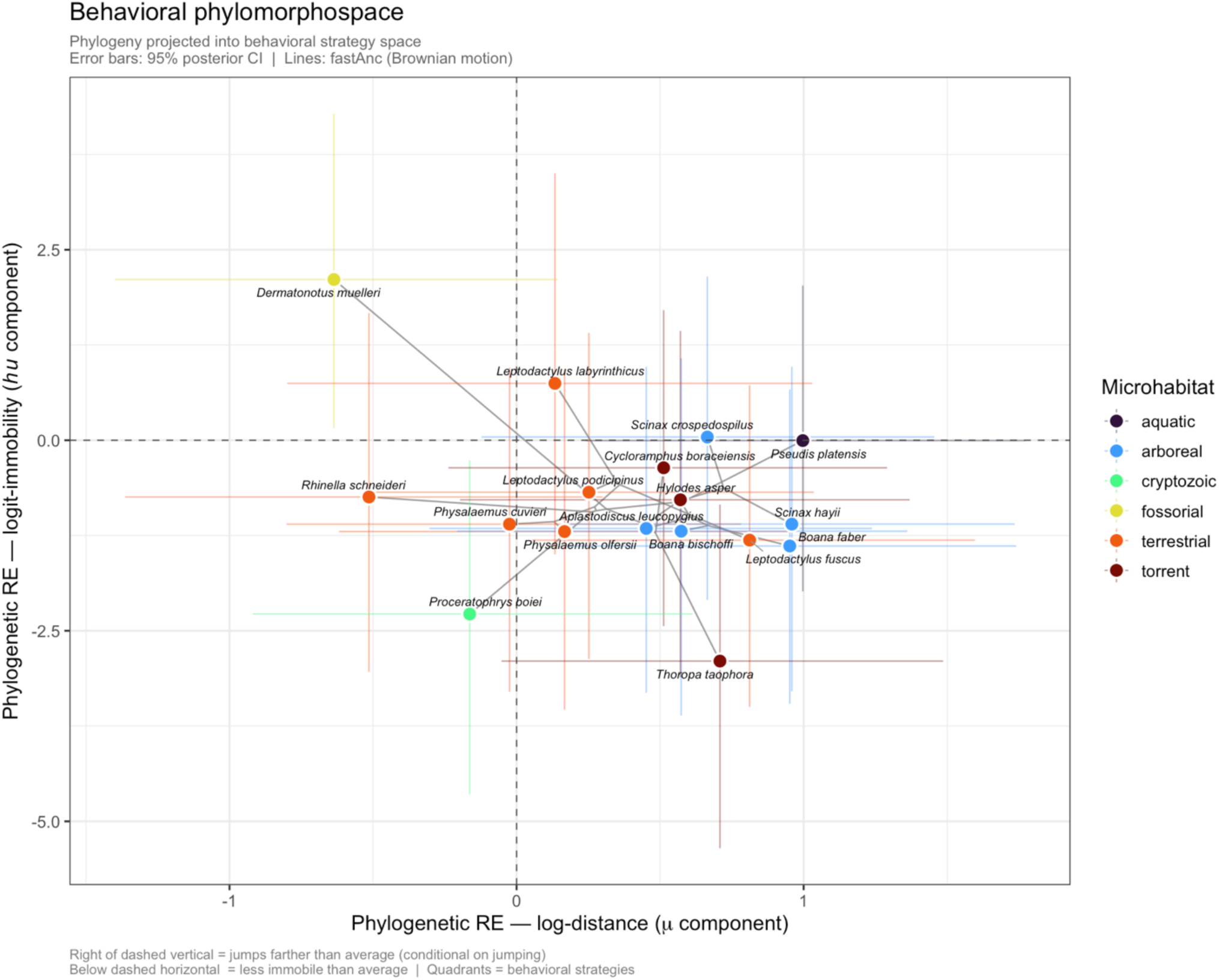
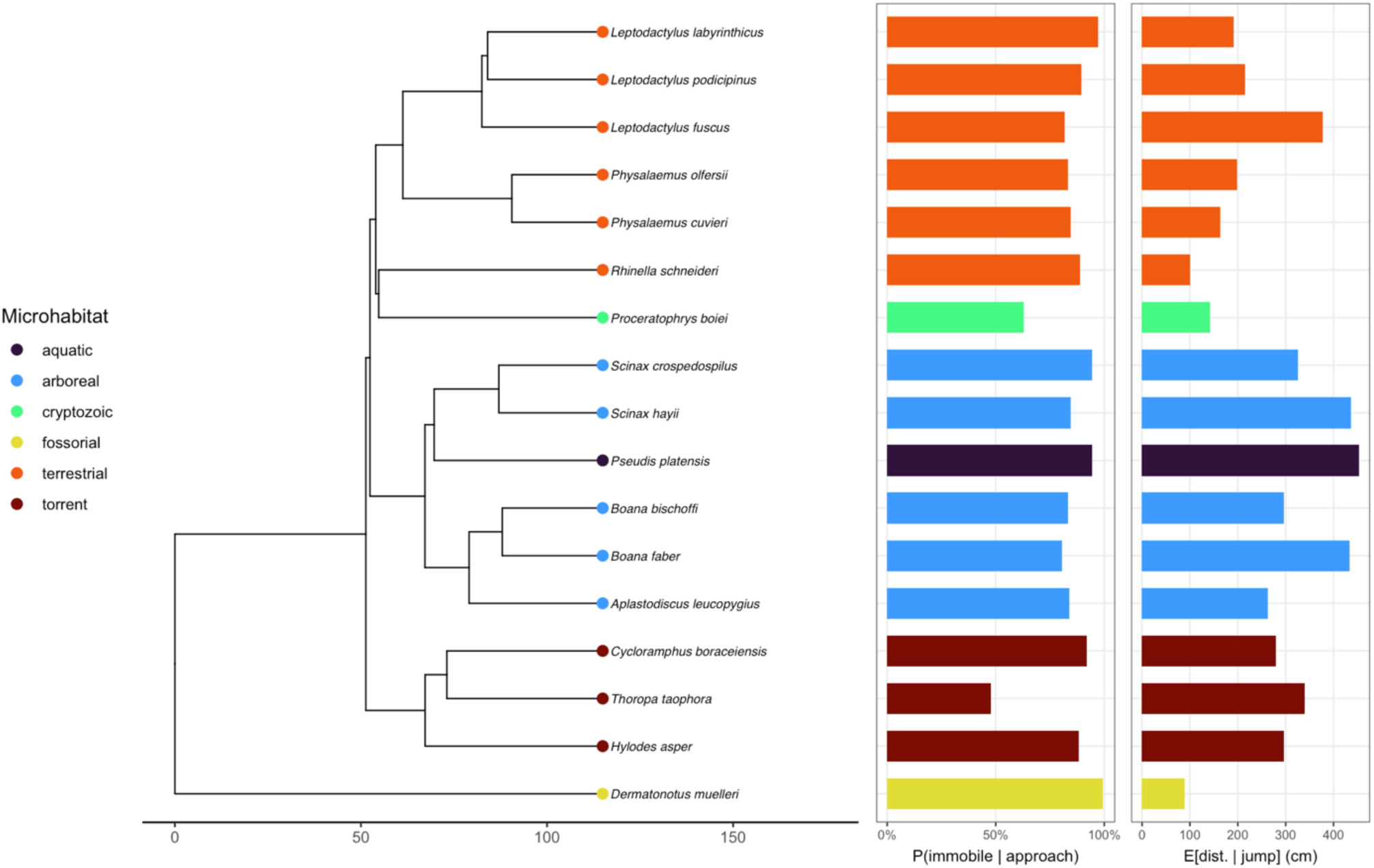
Behavioral phylomorphospace and phylogeny with estimated probabilities for the two model components. (a) Two-dimensional behavioral strategy space defined by posterior median species Random Effects: *mu* RE (x-axis: deviation in log-distance from the population mean) and *hu* RE (y-axis: deviation in logit-immobility). The phylogeny is projected onto the space with internal node positions estimated under a Brownian motion model. Error bars show posterior 95% ETIs. (b) Phylogeny showing P(immobile|approach) and E[distance|jump] for each species.

**Figure 5.**
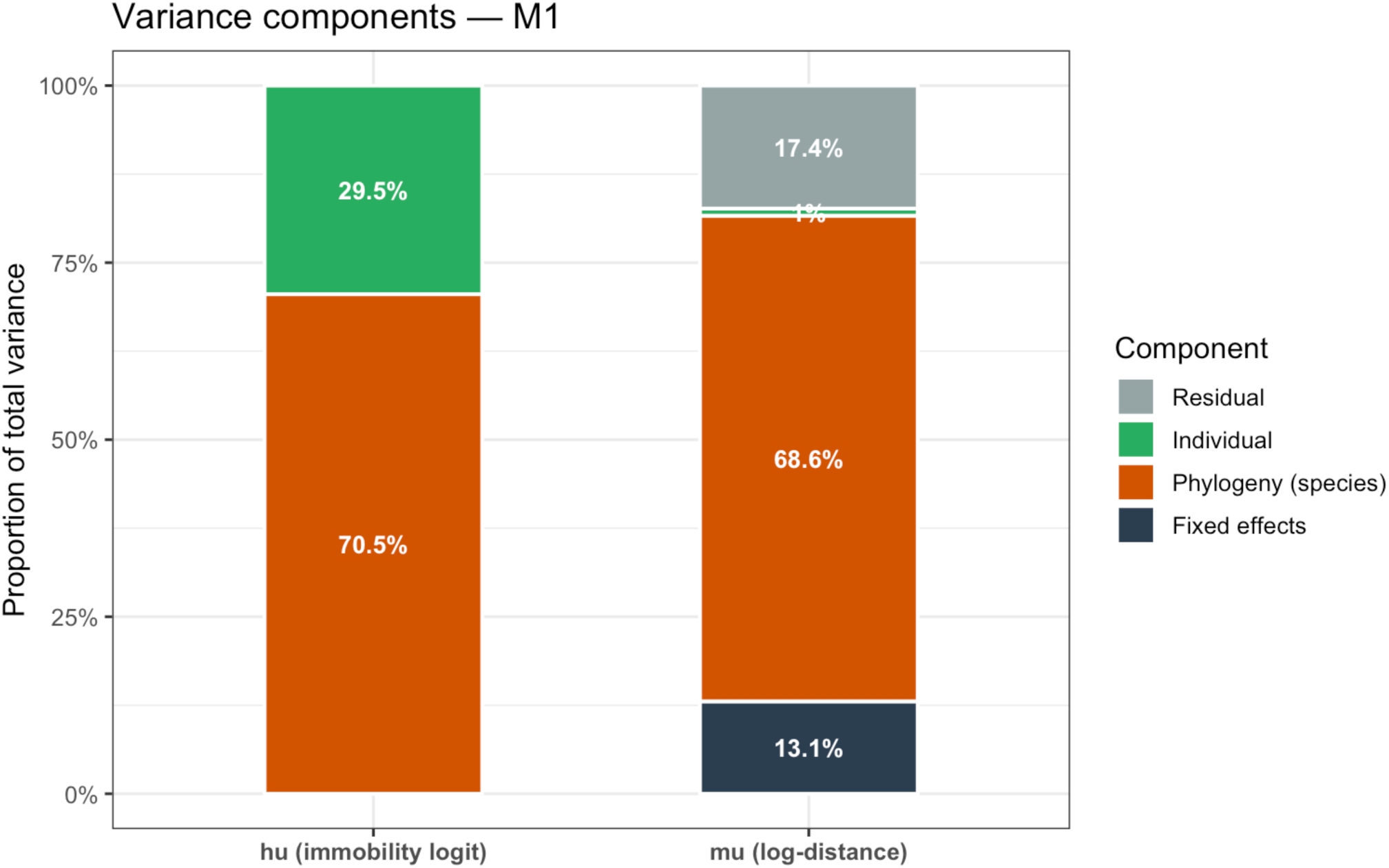
Variance partitioning of the two model components, showing the proportion of total variance explained by fixed effects, phylogenetic random effects, individual random effects, and residual variance for both the *mu* (conditional jump distance) and *hu* (immobility probability) components. Phylogenetic variance dominates both components (69.3% for *mu*; 75.1% for *hu*).

### Model comparison

LOO-CV confirmed reliable diagnostics for M1: all Pareto-*k* < 0.7 (Figure S8). M2 confirmed that the arena and stimulus effects on conditional jump distance are robust to the modelling choice: the equivalent parameters in M2 showed the same sign and similar magnitude as in M1, validating the hurdle formulation.

## DISCUSSION

Our Bayesian phylogenetic hurdle log-normal model of antipredator behavior in 17 Neotropical frog species revealed a consistent and biologically coherent picture: escape decisions are shaped by the immediate risk conveyed by predator stimuli, modulated by the physical opportunity that habitat structure provides for successful crypsis, and constrained by body size and deep evolutionary history. Predator touch, the clearest proximate cue of imminent capture risk, was the single most powerful predictor in the model, collapsing the probability of immobility from a near-certain baseline of 94% under visual approach to just 15% under physical contact, while simultaneously driving a 26% increase in conditional jump distance among individuals that escaped actively. These complementary shifts confirm that anurans switch seamlessly between cryptic and active escape strategies depending on the stage of the predation sequence (Gomes et al. 2002; Nishiumi and Mori 2015; Kikuchi et al. 2023), and that this switching carries a strong phylogenetic imprint (λ*_mu_* ≈ 0.80; λ*_hu_* ≈ 0.71).

Optimal escape theory (OET) predicts that prey should adopt the strategy that minimizes the sum of predation risk and opportunity cost of escape, and that the transition from passive to active defense should occur when risk exceeds a threshold set by the cost of fleeing (Ydenberg and Dill 1986; Cooper and Frederick 2007; Blumstein et al. 2016). Physical contact with a simulated predator represents the most direct signal that this threshold has been crossed. Consistent with this prediction, most individuals switched from immobility to active flight upon touch, regardless of arena type. This corroborates classic work on distance-dependent strategy switching in frogs (Nishiumi and Mori 2015) and aligns with the broader framework of predation sequences (Langerhans 2006; Kikuchi et al. 2023): pre-contact stages favor cryptic defenses that avoid detection, whereas post-contact stages favor active performance-based escape.

The complementary increase in jump distance under touch (+26%) reveals that individuals committed more forcefully to flight once they chose to escape. This is consistent with the all-or-nothing character of active escape decisions proposed by OET: once detection has occurred and predator contact is imminent, a partial escape that merely relocates the individual within striking distance confers little benefit (Cooper et al. 2015; Samia et al. 2016). Analogous results have been reported in frogs (Gomes et al. 2002), salamanders (Ducey and Brodie 1983), and in comparative studies of cryptic *vs*. aposematic frogs (Blanchette et al. 2017). Even mimetic frogs that benefit from bold behavioral signals initiate broader escape trajectories precisely when they face the risk of physical predator sampling (Barnett et al. 2025).

The higher immobility probability in the structurally complex bush arena relative to empty and leaf-litter arenas is consistent with the prediction that physical refuge availability and visual concealment increase the expected payoff of crypsis (Lima and Dill 1990). In complex environments, the probability that a motionless, camouflaged prey is detected decreases substantially (Wheatley et al. 2020; Fakan et al. 2023), making immobility a higher-payoff strategy than in open terrain, where the prey silhouette is fully exposed. Our results extend this logic across habitats: even the semi-structured leaf-litter treatment modestly increased immobility relative to the empty arena, suggesting that the benefit of complexity operates on a continuous scale rather than as an all-or-nothing switch.

The simultaneous increase in conditional jump distance in open arenas (+31% in both empty and leaf-litter relative to bush; pd ≥ 99.7% for both) is precisely what OET predicts when refuges are absent or distant. In the absence of nearby cover, a short jump merely relocates the prey without reaching safety, so the optimal active escape requires a longer displacement (Martín et al. 2005). Iberian green frogs jumped shorter distances when aquatic vegetation was present (Martín et al. 2005); tree frogs jumped shorter distances in complex natural habitats than in open laboratory arenas (Gomes et al. 2002). In southern leopard frogs, open microhabitats delayed flight relative to grass cover, and the effect of body size was strongest in vegetated microhabitats (Bateman and Fleming 2014). A broader multi-species field study confirmed that species differ sharply in the balance between freezing and fleeing across open and covered microhabitats, with small species becoming more mobile in the open while larger ground-dwelling remained still (Matich and Schalk 2019). Our study replicates and generalizes these patterns across 17 species and six families in a controlled comparative framework, providing the first multi-species phylogenetic estimate of the arena effect on both escape strategy and jump distance. The arena × stimulus interactions on conditional distance were near zero (pd < 58% for empty × touch; pd = 91.8% for leaf litter × touch), indicating that these effects are approximately additive on the log scale. This additivity is consistent with an economic model in which stimulus type determines the intensity of escape commitment and habitat structure independently determines the displacement required to reach safety (Ydenberg and Dill 1986; Cooper and Blumstein 2015).

The effect of body size on immobility probability was among the clearest signals in the model: a one-SD increase in log SVL increased the logit of immobility by 2.37 ± 0.71 units (pd > 99.9%), shifting P(immobile) from ∼77% to ∼96% at the population-mean intercept. By contrast, the same covariate showed only a weak positive tendency on conditional jump distance (β = +0.246 ± 0.182; pd = 91.3%). Taken together, body size is primarily a determinant of which strategy to use rather than how intensely to execute active escape. This dissociation between strategy-selection and locomotor-performance roles of body size is consistent with findings across taxa. Larger frog species tend to have greater limb muscle mass, but lower size-corrected jumping power (Mendoza et al. 2020), such that large body size does not automatically confer superior active escape performance. As body mass increases, jumping power decreases, with the fastest decline in burrowing species (Mendoza et al. 2020), a trade-off directly relevant to *Dermatonotus muelleri*, whose fossorial body plan — characterized by short hindlimbs and a pelvic morphology associated with burrowing locomotor mode (Jorgensen and Reilly 2013) — may simultaneously constrain jumping power while relaxing selection pressure on active escape. In chameleons, body size similarly predicts which antipredator strategy is adopted rather than the speed of any given strategy (Drown et al. 2022). The principle extends beyond body size to other axes of locomotor capacity: in dice snakes (*Natrix tessellata*), individuals with intrinsically slower locomotion resort proportionally more to death feigning (Bjelica et al. 2024). Across these cases, organisms appear to shift toward defenses that do not depend on locomotor performance whenever that performance is constrained: whether by body plan, by body size, or by individual variation alike. Importantly, body size also shapes which defensive channels are available. For example, the probability of possessing distress calls rises steeply above ∼52 mm SVL (Forti et al. 2018), suggesting a size threshold for certain acoustic defenses. Among chemically defended frogs, conspicuous coloration is more likely in smaller diurnal species (Roberts et al. 2022). Body size thus changes defense architecture, not merely locomotor capacity, making it plausible that larger species rely more on immobility without necessarily jumping farther once they do flee.

Our estimates — λ*_mu_* ≈ 0.80 [0.70, 0.94] for conditional jump distance and λ*_hu_* ≈ 0.71 [0.55, 0.90] for immobility probability — indicate moderately strong phylogenetic conservatism in both response dimensions. Pagel’s λ quantifies the degree to which trait variation is structured by phylogenetic relatedness relative to a Brownian motion model (Pagel 1999; Pearse et al. 2025). These values are remarkably concordant with phylogenetic signal estimates for anuran locomotor performance (Mendoza and Moen 2020), in which multivariate signal for anatomical approximations of jumping performance at K_mult_ = 0.71. The convergence between behavioral signal estimates (λ ∼ 0.7–0.8) and performance signal estimates (K ∼ 0.7–0.8) across independent datasets and methods suggests that the evolutionary dynamics of jumping behavior and jumping morphology are coupled.

The phylogenetic variance component dominated the total variance in both the *mu* component (69.3%) and the *hu* component (75.1%). These high proportions indicate that most interspecific behavioral variation reflects deep evolutionary divergences among lineages. Frog microhabitat diversification drove evolutionary changes in hindlimb morphology consistent with multiple Ornstein–Uhlenbeck optima (Citadini et al. 2018, Buttimer et al. 2020), and earlier work demonstrated that pelvic and hindlimb skeletal variation is phylogenetically structured and associated with locomotor mode (Reilly and Jorgensen 2011; Jorgensen and Reilly 2013; Leavey et al. 2023). Our results suggest that antipredator behavioral strategy has co-diversified with morphology across the same microhabitat gradients. Importantly, the strong signal we report is consistent with recent clade-level comparative studies. In *Proceratophrys* (Odontophrynidae) — a genus represented in our sample by *P. boiei* — Zocca et al. (2022) found substantial phylogenetic structure in camouflage, immobility, distress calls, and aggression, with camouflage and immobility being particularly conserved. Crouching, body inflation, rear elevation, and eye protection evolved in Leiuperinae (Leptodactylidae) as linked syndromes rather than as independent traits (Ferraro et al. 2020), suggesting that the strong signal in our immobility component may partly reflect the conserved nature of integrated defensive sequences. Ferreira et al. (2019) further showed that although the broad defense repertoire is ancestral and widespread, individual traits, such as distress calls are convergent, indicating that strong phylogenetic signal and ecological tuning are not mutually exclusive.

The behavioral phylomorphospace organized species into a recognizable ecological structure. Fossorial *Dermatonotus muelleri* occupied the extreme positive end of the immobility axis. Arboreal species consistently occupied the lower half of the immobility axis, indicating greater propensity for active flight, consistent with the ecomorphological prediction that arboreal frogs have longer hindlimbs and higher jumping performance (Citadini et al. 2018; Buttimer et al. 2020). The torrent specialist *Thoropa taophora* occupied the active-escape quadrant, consistent with the demanding locomotor environment of rocky torrent margins. The clustering of closely related species within shared behavioral quadrants reinforces the phylogenetically structured nature of strategy differences.

The individual-level variance for the immobility component (SD_ind,hu_ = 1.145) was substantial, accounting for approximately 24.9% of total *hu* variance. This means that, above and beyond species-level and phylogenetic effects, individual frogs of the same species consistently differ in their immobility tendency across repeated trials. Such repeatability of individual behavior is a necessary condition for individual differences to be targeted by selection (Dingemanse and Dochtermann 2013). Formalizing this as within-species adjusted repeatability (Nakagawa and Schielzeth 2010; Stoffel et al. 2017), we estimated R_hu_ = 0.30 (95% ETI: [0.12, 0.49]), indicating that ∼30% of within-species variation in immobility tendency is attributable to consistent among-individual differences. Within-species individual variation in antipredator responses has been documented in frogs (Castellano et al. 2023), salamanders (Waldron et al. 2021), and toads (Chiocchio et al. 2022; Baskiera et al. 2021). By contrast, repeatability of conditional jump distance was low (R_mu_ = 0.07; 95% ETI: [0.00, 0.19]), confirming that conditional jump distance is more environmentally determined than individually consistent. Individual frogs are thus consistent in *which* strategy they deploy, but not in *how far* they jump when they flee, a pattern consistent with the view that strategy choice and locomotor intensity reflect functionally distinct levels of behavioral organization (Arnold 1983; Garland and Losos 1994).

This study has some limitations. First, only males were used, which precludes inference about sexual differences in antipredator strategy (Zamora-Camacho et al. 2022). Second, our sample includes 17 species from the Brazilian Atlantic Forest, which does not represent the full phylogenetic breadth of frogs. Third, our arenas were standardized experimental conditions rather than natural microhabitats. Fourth, the model does not include information on natural predator community composition, which varies substantially across species ranges. Incorporating predator diversity or predation pressure indices as additional covariates could help partition the phylogenetic random effects into historical and contemporary ecological components.

## Conclusion

We demonstrate that anuran antipredator behavioral strategies: the choice between immobility and active flight, and the intensity of flight when it occurs are shaped by the interaction of predator stimulus type, habitat structural complexity, body size, and evolutionary history. Predator contact acts as a near-universal switch from crypsis to flight across species, habitat contexts, and body sizes. Habitat complexity amplifies the benefit of crypsis and reduces the required escape distance. Body size primarily determines which strategy is optimal rather than how powerfully active escape is executed. The strong phylogenetic signal in both decision-making and locomotor response components suggests that the evolutionary history of microhabitat use has co-shaped both morphological and behavioral diversity, and that escape decisions are deployed from broader, phylogenetically conserved defense syndromes rather than assembled anew in each species (Ferreira et al. 2019; Zocca et al. 2022; Ferraro et al. 2020). The substantial individual variation in immobility tendency provides a heritable substrate on which ongoing natural selection could act within populations.

## USE OF GENERATIVE AI TOOLS

In accordance with the guidelines of Oxford University Press, COPE (2023), and following the AIdIT framework (Drobniak et al. 2026), we disclose the following use of generative artificial intelligence tools during the preparation of this manuscript.

### AI tool(s) used

Claude (Anthropic, models Sonnet 4 and Opus 4).

### Area(s) of generative AI usage (AIdIT taxonomy)

- *Writing — text editing for style/grammar/spelling:* AI was used to assist with language editing, improving sentence structure and clarity in selected sections of the manuscript.
- *Writing — text generation based on prompts:* AI was used to produce initial drafts of selected manuscript sections (Discussion, Results) from structured prompts containing detailed statistical outputs, hypotheses, and author-provided interpretive notes. All generated text was critically reviewed, extensively revised, and rewritten by the authors to ensure accuracy and consistency with the data and the authors’ interpretations.
- *Formal analysis — refining existing code:* AI was used to debug and optimize R code for Bayesian model fitting (brms), posterior processing, and figure generation. All analytical decisions, model specifications, prior choices, and interpretations were made by the authors.
- *Visualisation — code generation for visuals:* AI assisted in generating R/ggplot2 code for data visualization. All figures were verified and approved by the authors.

### Human oversight statement

The authors declare that they have verified and approved all content generated or modified by the AI tools used. No AI tool was used for data collection, experimental design, statistical model selection, or interpretation of results. The AI tool is not listed as an author because it does not meet authorship criteria (ICMJE/COPE). The authors take full responsibility for the content of this publication, including any parts informed by generative AI tools.

## ACKNOWLEDGMENTS

This manuscript is part of JMC PhD dissertation, as such she would like to thank Carlos A. Navas and Cinthia A. Brasileiro as members of her dissertation examination committee. Diogo S. M. Samia commented on an earlier draft of this manuscript and provided useful ideas and guidance. Maya Maia, Adriana Barsotti, Eduardo Moretti, Inês Rosa, Faride, Camila, Jesus, Luciano A. Anjos, Juma, Otilie, Douglas, Murilo, Cristiele, Fábio P. de Sá, and several members of the Laboratory of Behavior and Evolutionary Physiology (IBUSP/SP) and Laboratory of Eco-physiology and Evolutionary Physiology (IBUSP/SP) helped with field work. The Butantan Institute (SP) provided JMC with the snakes used in the experiment.

## AUTHOR CONTRIBUTIONS (CRediT)

**Diogo B. Provete:** Formal analysis, Methodology (contributing), Software, Visualization, Writing – original draft (lead), Writing – review & editing (lead). **Jessyca M. Citadini:** Conceptualization (equal), Investigation (lead), Methodology (equal), Data curation, Writing – original draft. **Fernando R. Gomes:** Conceptualization (equal), Funding acquisition, Methodology (equal), Resources, Supervision, Writing – review & editing (equal).

## DATA AVAILABILITY

Analyses reported in this article can be reproduced using the data and the Quarto documents provided by Anonymous (2026).

## SUPPLEMENTARY MATERIAL

### Behavioral Ecology

This document contains the supplementary material for the manuscript “Predator stimulus and habitat structure jointly shape antipredator behaviour in frog species”. It includes **Table S1** (species-level sample sizes and microhabitat classification), **Figure S1** shows the map with sampling localities and **Figure S2** (photographs of the three experimental arenas). **Figures S3–S6** present MCMC diagnostics — trace plots, posterior- and prior-predictive checks, and prior sensitivity analysis via the R package *priorsense* — confirming convergence and robustness of Model M1. **Figures S7–S9** report the posterior density of Pagel’s λ for both response components, Pareto-*k* leave-one-out cross-validation (LOOCV) diagnostics, and probability-of-direction (*pd*) and Region of Practical Equivalence (ROPE) analyses for the fixed-effect coefficients. **Figures S10–S12** describe the empirical distribution of jump distance, the proportion of zero observations (immobility) by species, and species-level random effects for both model components. **Figures S13 and S14** present marginal effects of body size and the arena × stimulus interaction on the linear predictors.

### TABLE S1

**Table S1.**
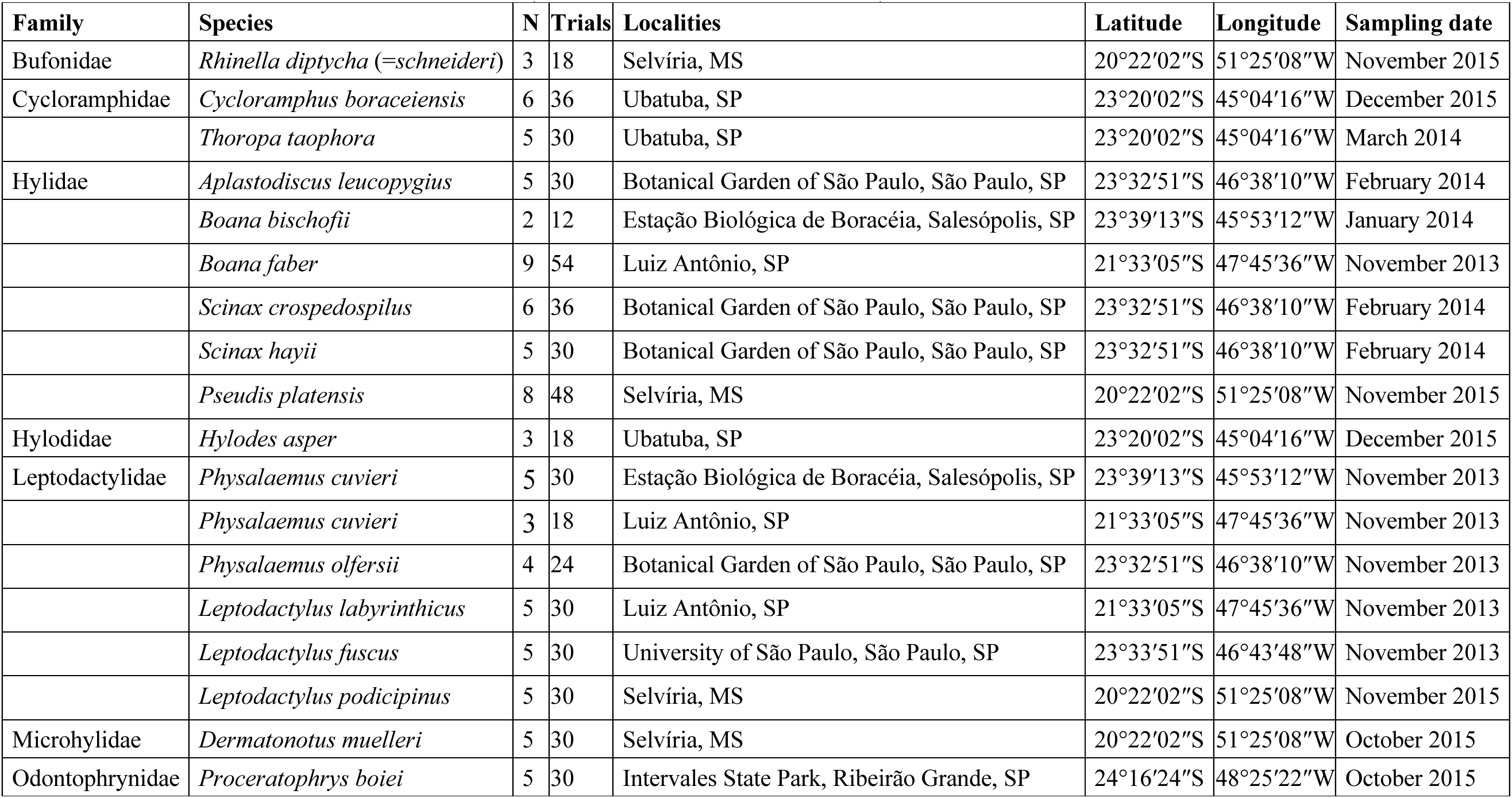
Species used in this study with sample sizes, sampling localities, geographic coordinates, and sampling dates. N = number of males tested; trials = total number of trials (N × 6 treatment combinations).

### FIGURES

**Figure S1.**
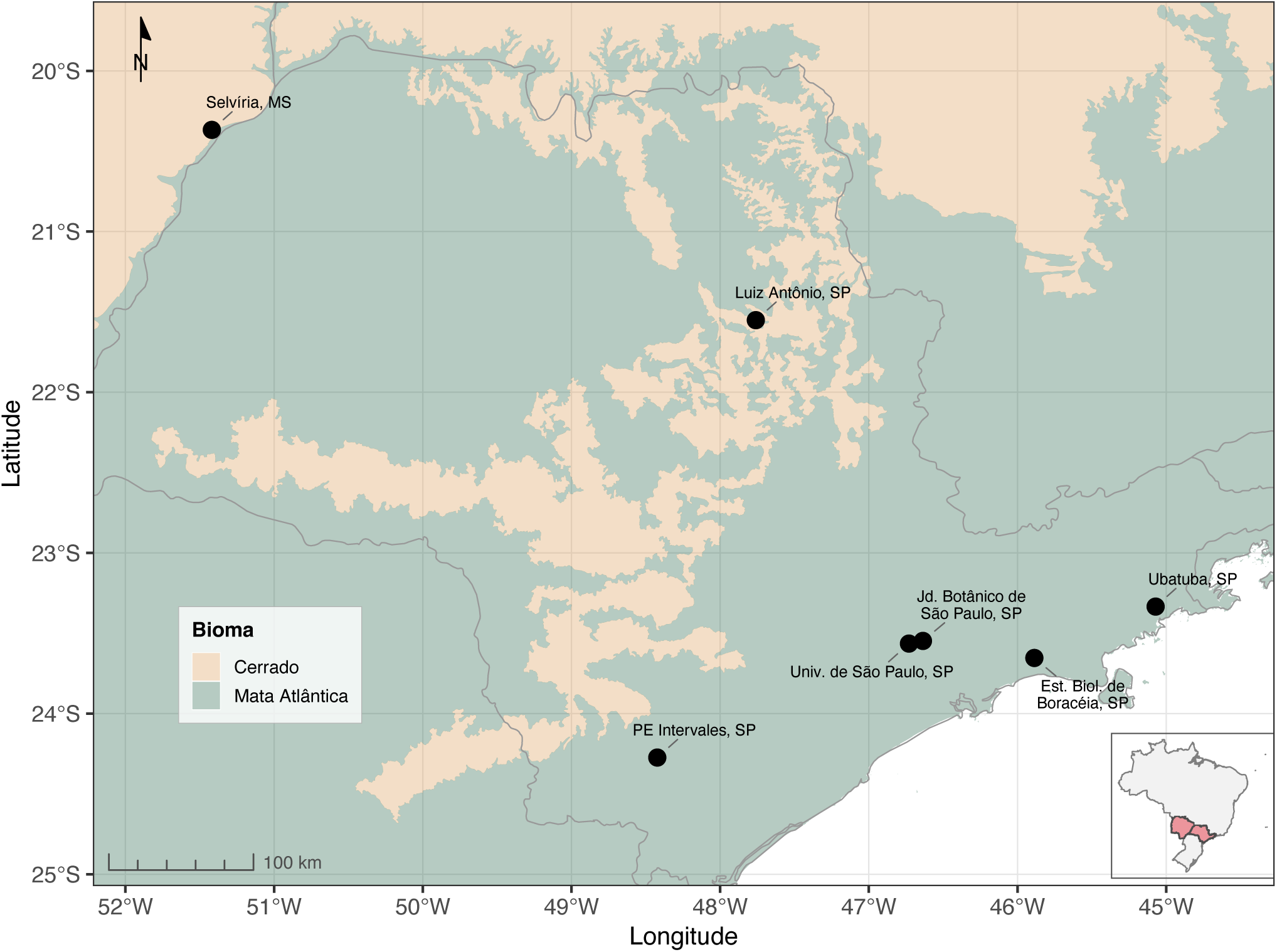
Sampling localities of the 17 anuran species used in this study. Black circles indicate the seven collection sites across the states of São Paulo and Mato Grosso do Sul, southeastern Brazil. Shaded areas represent the two biomes encompassed by our sampling: Cerrado (savanna-like vegetation in the interior) and Atlantic Forest (coastal and montane forests). Biome boundaries follow the Brazilian Institute of Geography and Statistics (IBGE, 2019). The inset shows the location of the study area within Brazil. See Table S1 for species, sample sizes, and coordinates of each locality.

**Figure S2.**
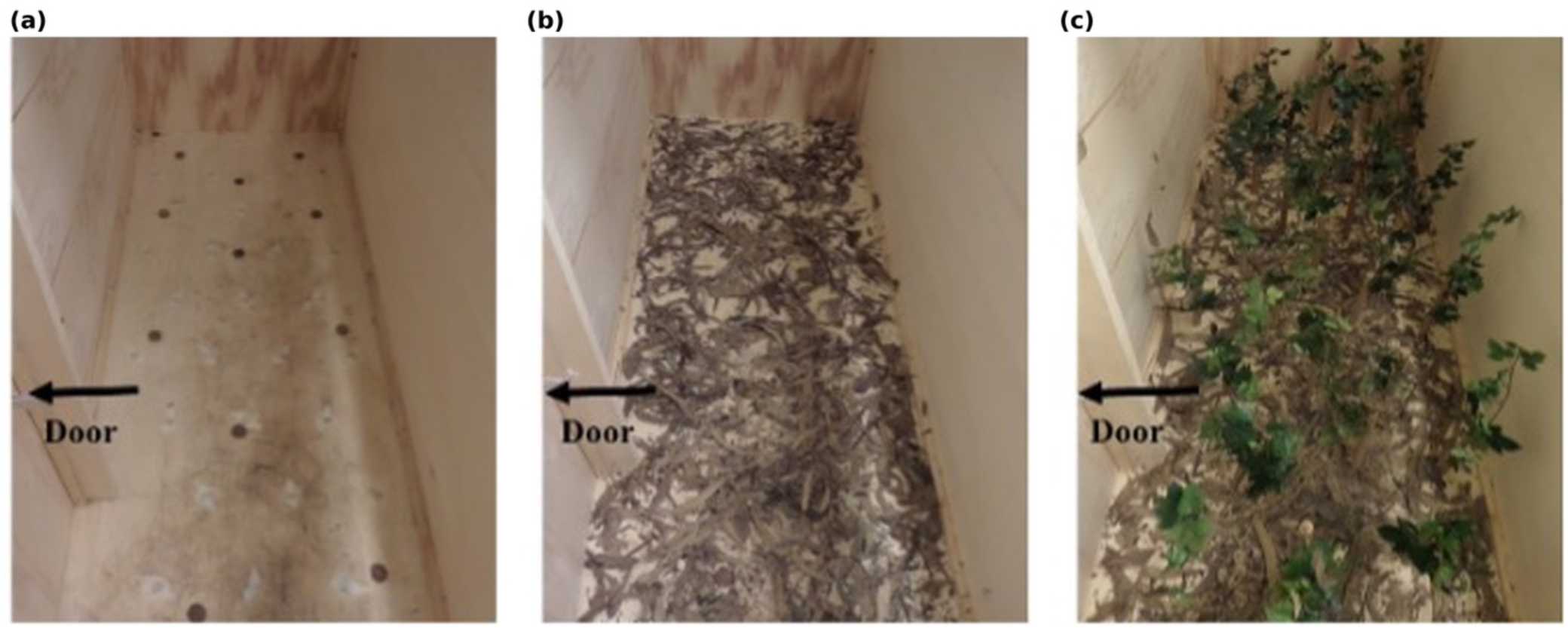
Pictures of the experimental arena (200 × 90 × 70 cm) showing the three structural complexity treatments: (a) empty arena, (b) leaf litter covering the floor, and (c) leaf litter with plastic bushes (50 cm height, spaced 10 cm apart). The small lateral door for frog placement is shown on the left side. The snake was introduced from the posterior end relative to the frog’s position.

### SUPPLEMENTARY FIGURES: MCMC DIAGNOSTICS

**Figure S3.**
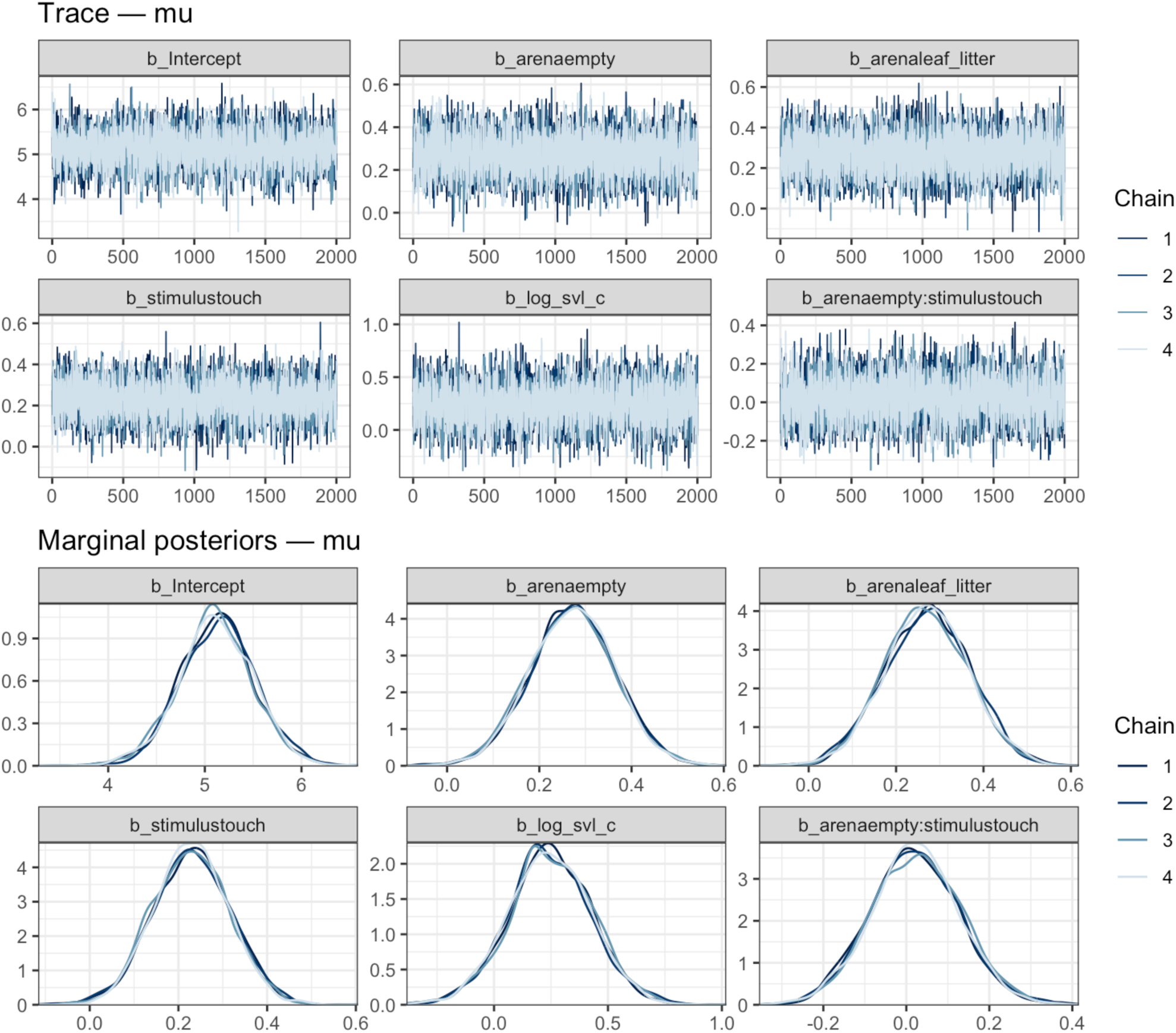
MCMC trace plots for all parameters of the mu (conditional jump distance) component of Model M1. Four parallel chains (6,000 iterations each, warm-up = 2,000, thinning = 2) are shown in different colors. All chains show good mixing with no systematic trends or stuck chains.

**Figure S4.**
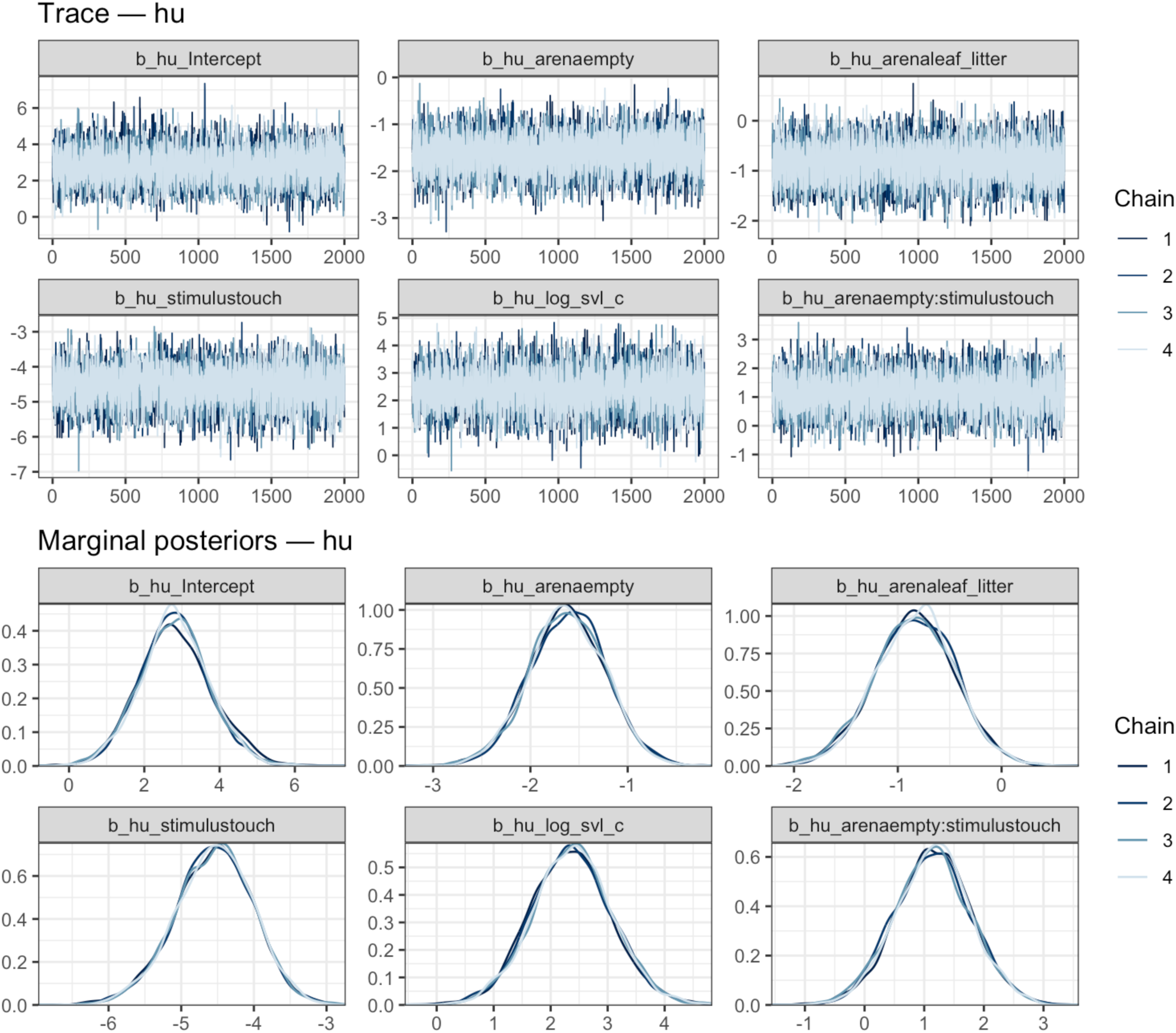
MCMC trace plots for all parameters of the hu (immobility probability) component of Model M1.

### SUPPLEMENTARY FIGURES: POSTERIOR PREDICTIVE CHECKS

**Figure S5a.**
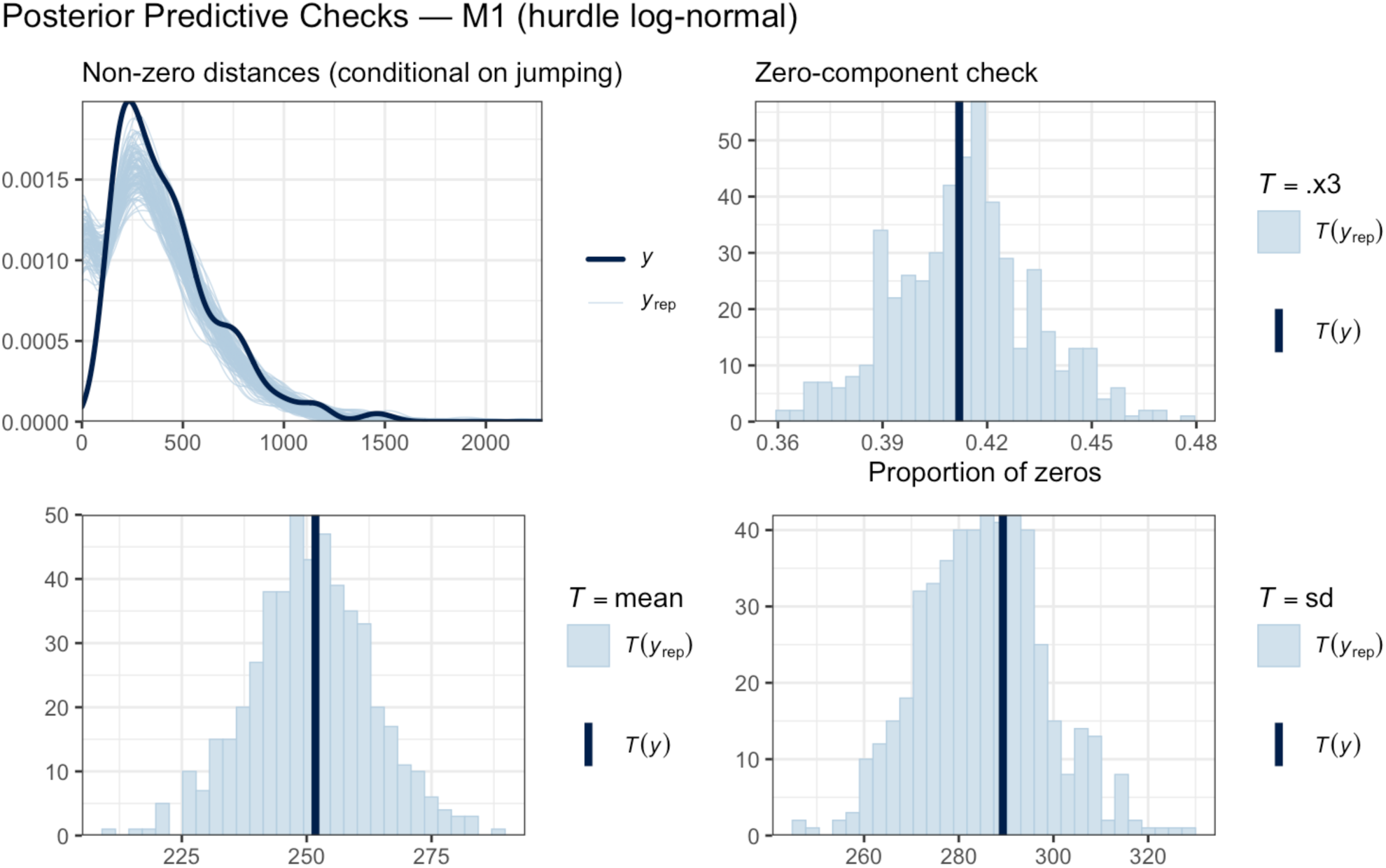
Posterior predictive check for Model M1 (hurdle log-normal, N = 534 trials). Dark blue line: observed data distribution; light blue lines: 100 replicated datasets drawn from the posterior predictive distribution. The model adequately recovers the observed proportion of structural zeros (∼41%) and the right-skewed conditional distribution of non-zero jump distances.

**Figure S5b.**
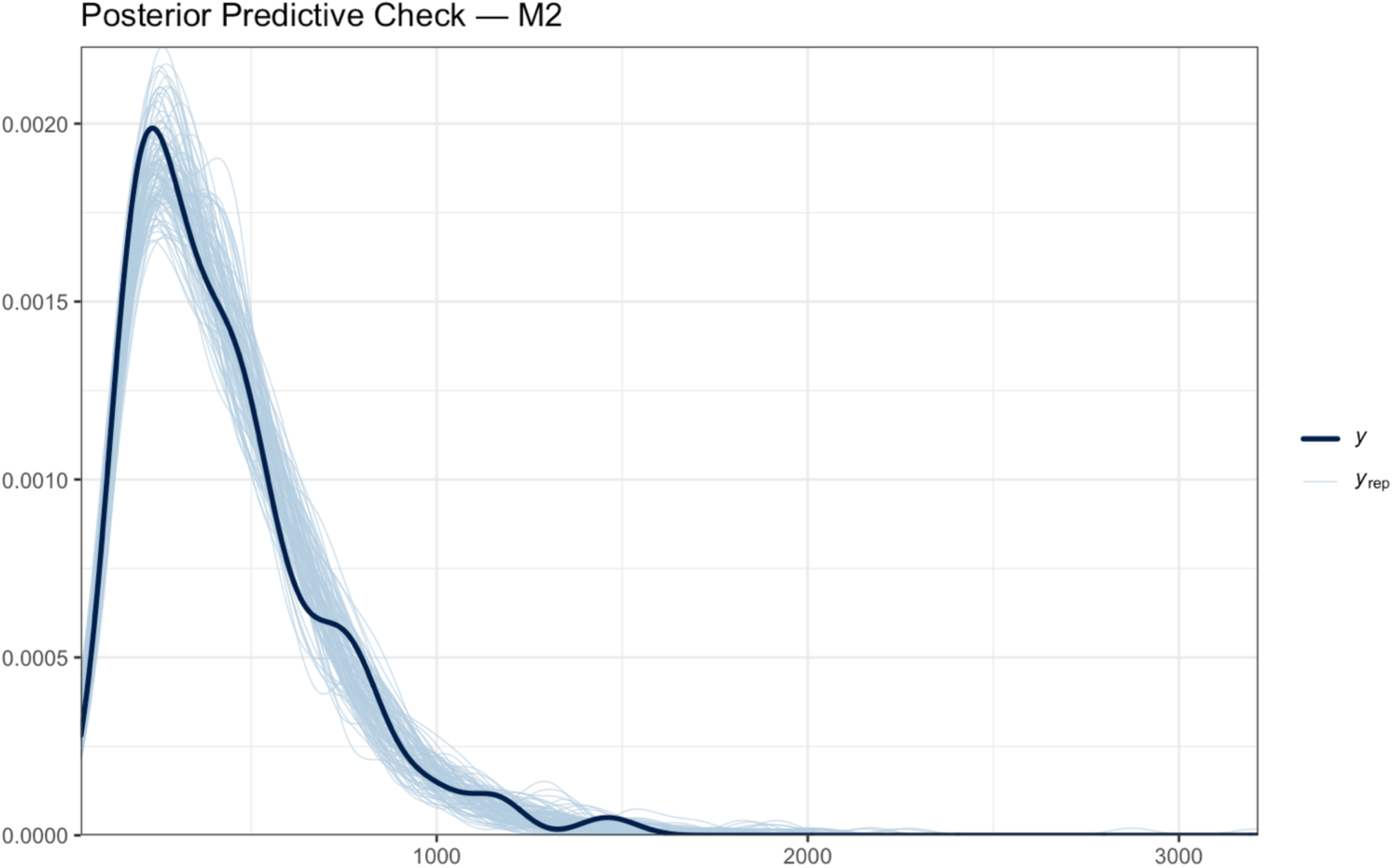
Posterior predictive check for Model M2 (log-normal, restricted to N = 314 non-zero observations).

**Figure S5c.**
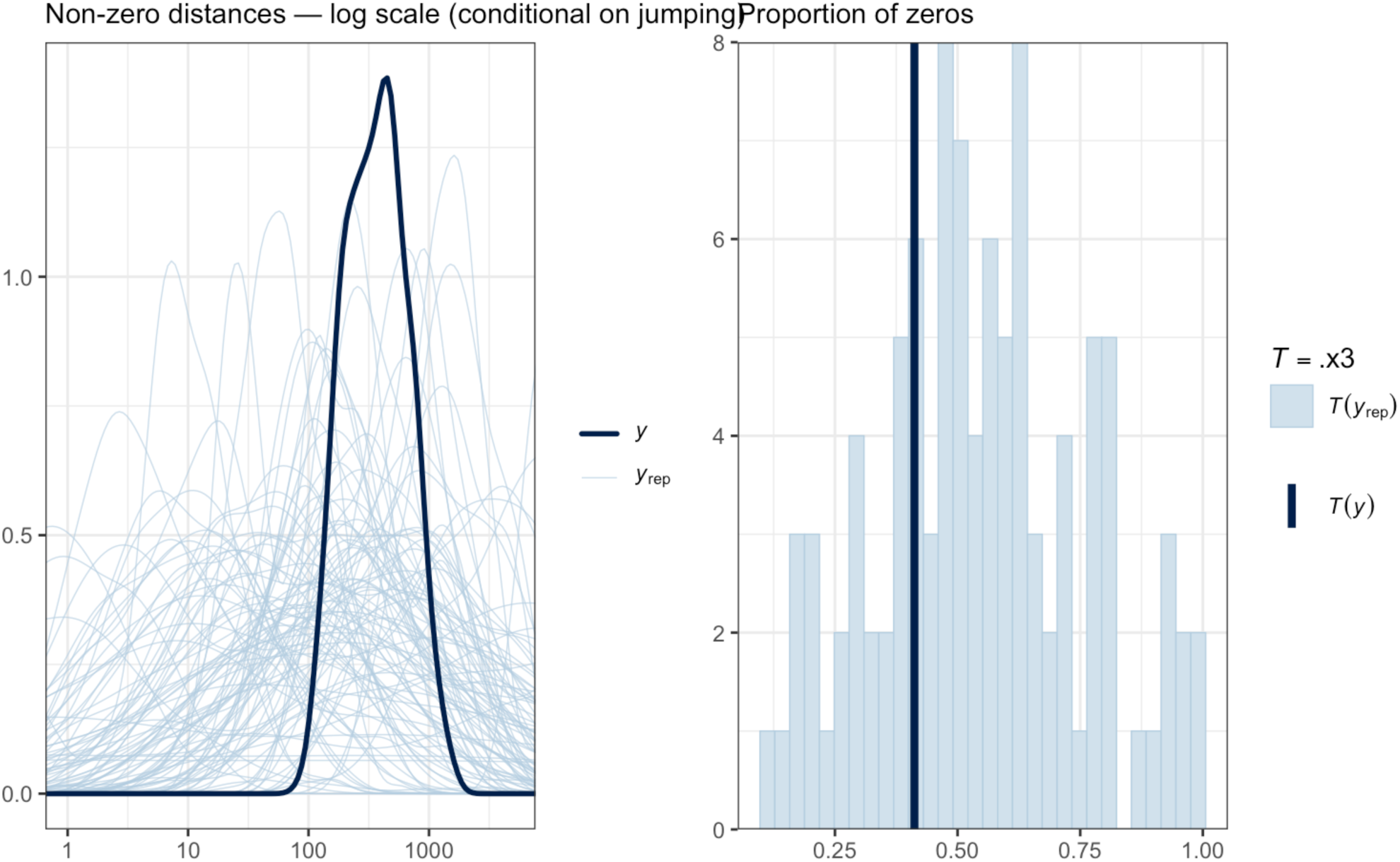
Prior predictive check. Simulated datasets generated from the prior distributions alone (before observing data). The prior generates plausible zero proportions (10–90%) and log-scale distances consistent with the observed range, confirming that priors are weakly informative, but not dogmatic.

### SUPPLEMENTARY FIGURES: PRIOR SENSITIVITY ANALYSIS

**Figure S6.**
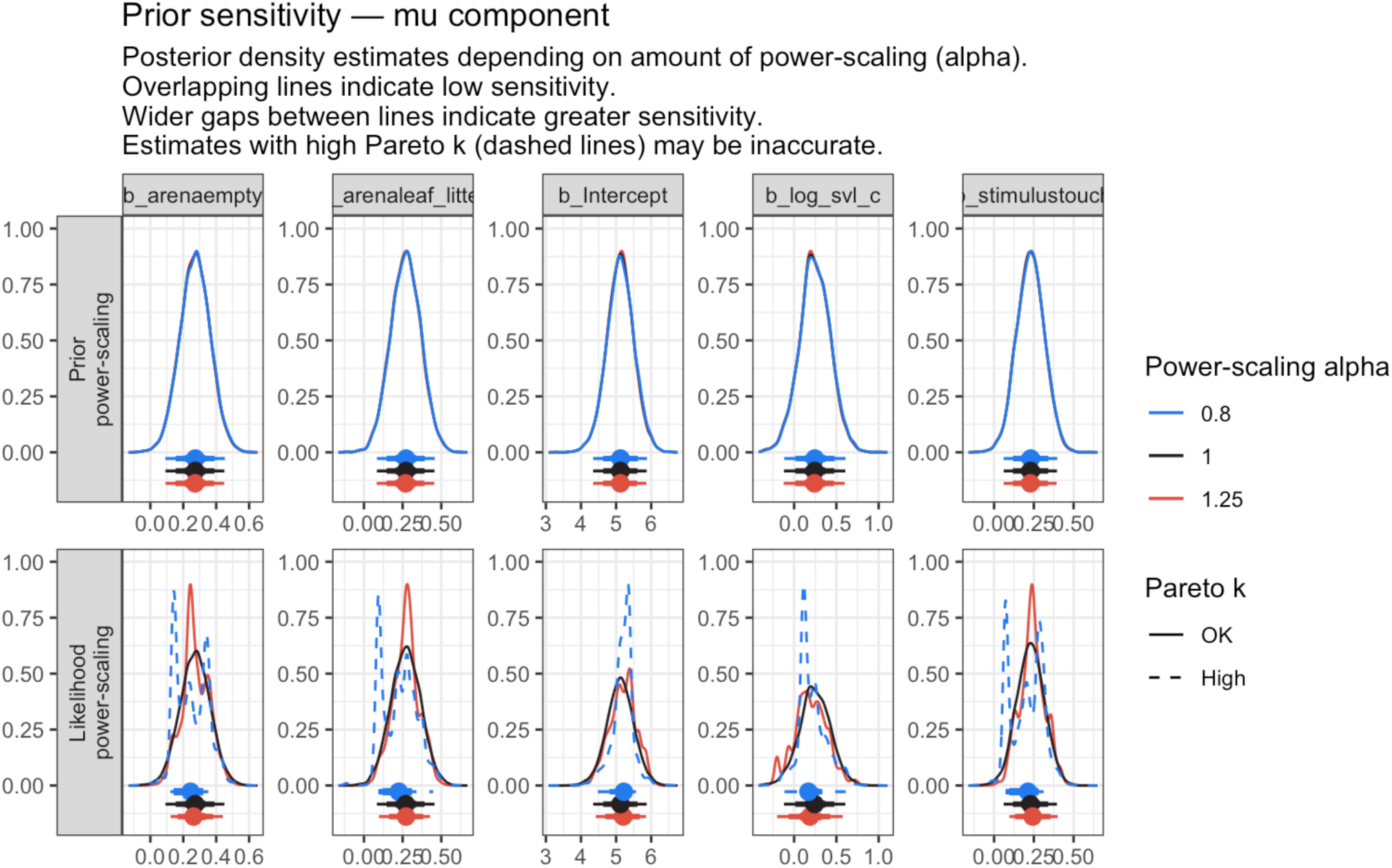
Prior sensitivity analysis for Model M1, implemented via the priorsense package (Kallioinen et al. 2024). Power-scaling diagnostic plots for all parameters. Posteriors were insensitive to ±25% prior perturbations for all *mu* parameters. Likelihood-scaling flags for some *hu* parameters reflect strong data informativeness — a desirable property confirming that the likelihood, not the prior, drives inference for these effects.

### SUPPLEMENTARY FIGURES: PHYLOGENETIC SIGNAL

**Figure S7.**
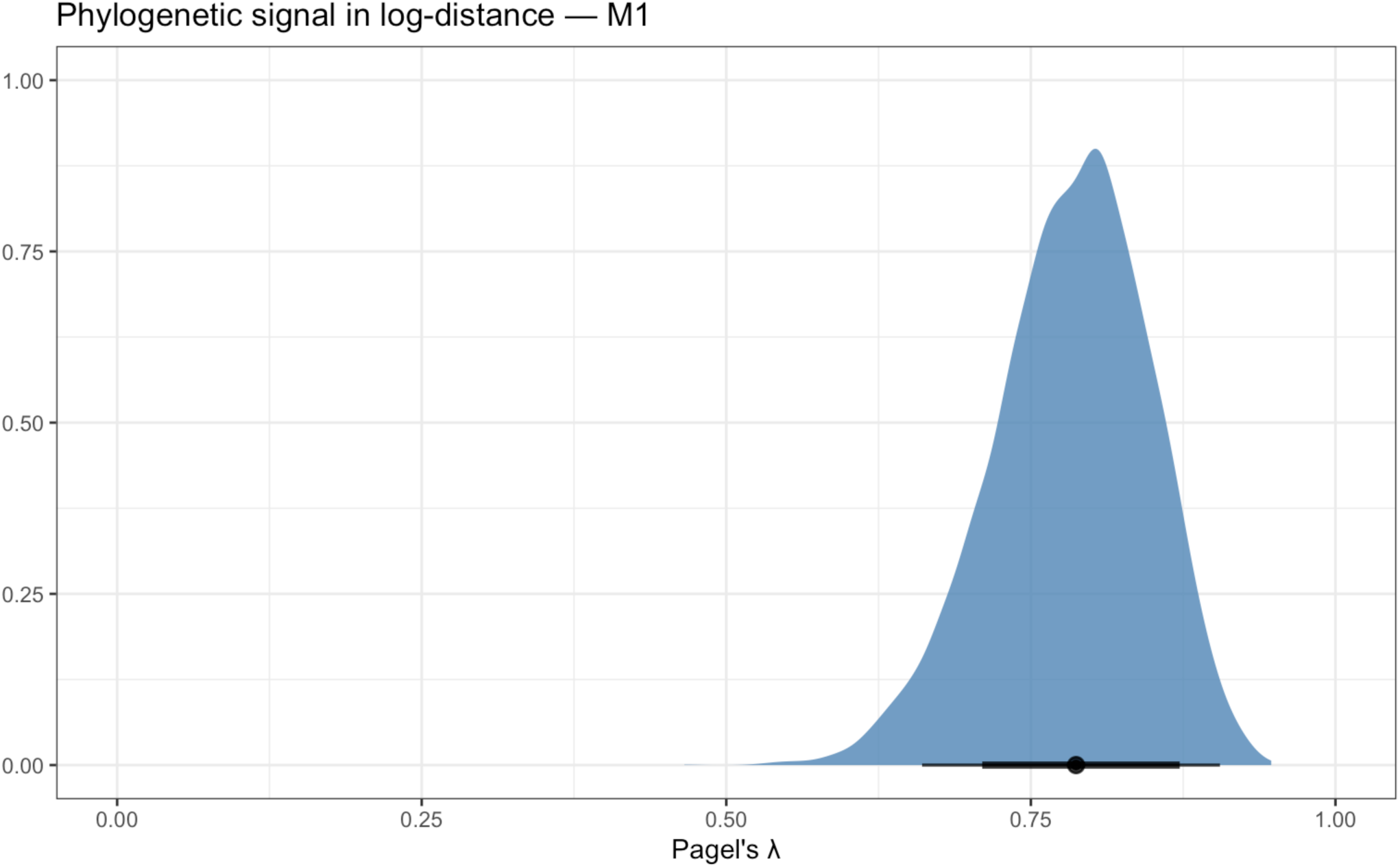
Posterior density of Pagel’s λ for both model components. λ*_mu_* ≈ 0.80 (95% Equal-Tailed Interval (ETI): [0.70, 0.94]) for conditional jump distance; λ*_hu_* ≈ 0.71 (95% ETI: [0.55, 0.90]) for immobility probability. Dashed vertical lines mark the posterior median; shaded area shows the 95% ETI.

### SUPPLEMENTARY FIGURES: LOO-CV DIAGNOSTICS

**Figure S8.**
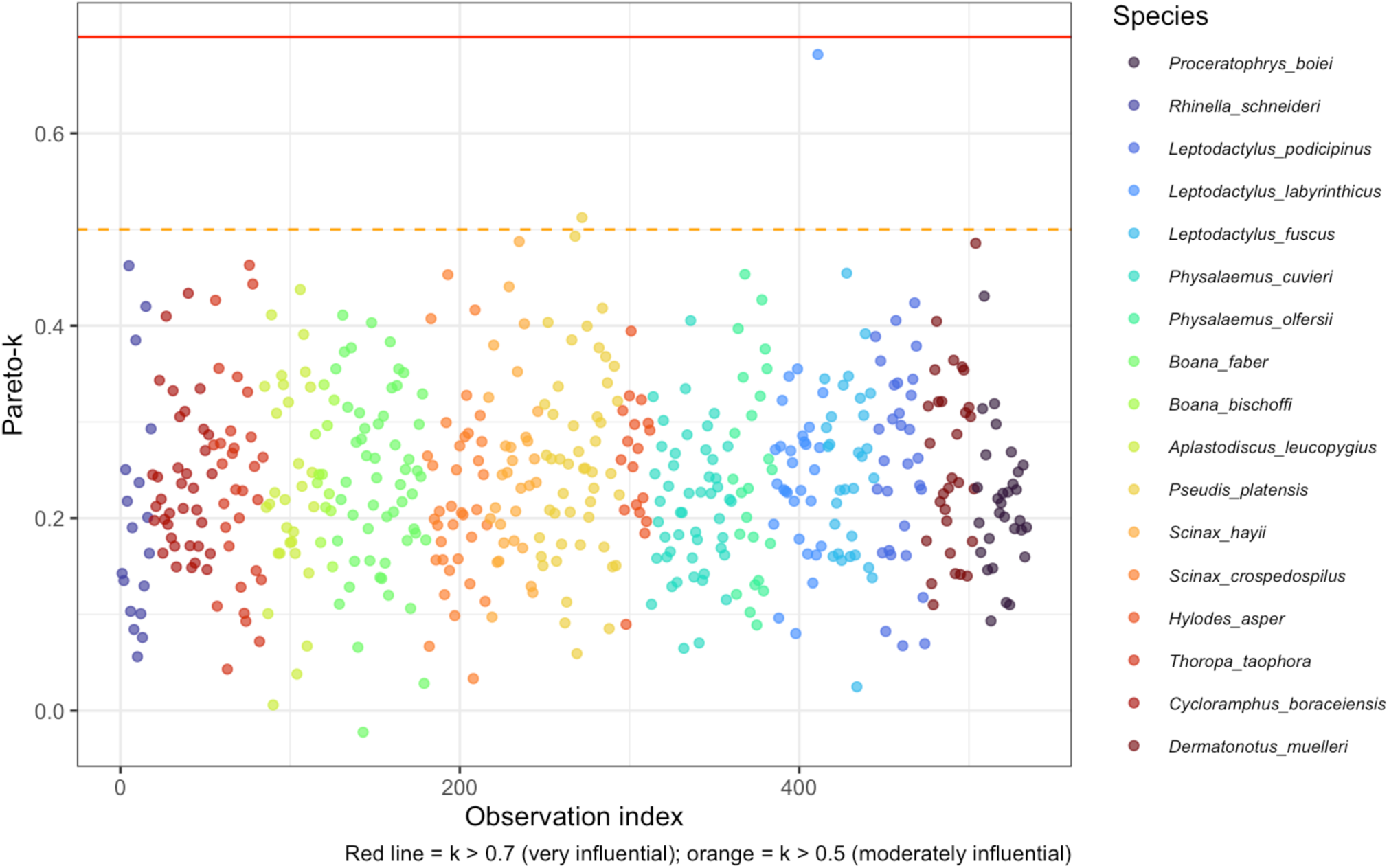
Pareto-*k* diagnostic plot for leave-one-out cross-validation (LOO-CV) of Model M1. All Pareto-*k* values < 0.7, confirming reliable LOO estimates. Points above 0.7 would indicate influential observations requiring moment-matching or refit.

### SUPPLEMENTARY FIGURES: ADDITIONAL DIAGNOSTICS

**Figure S9.**
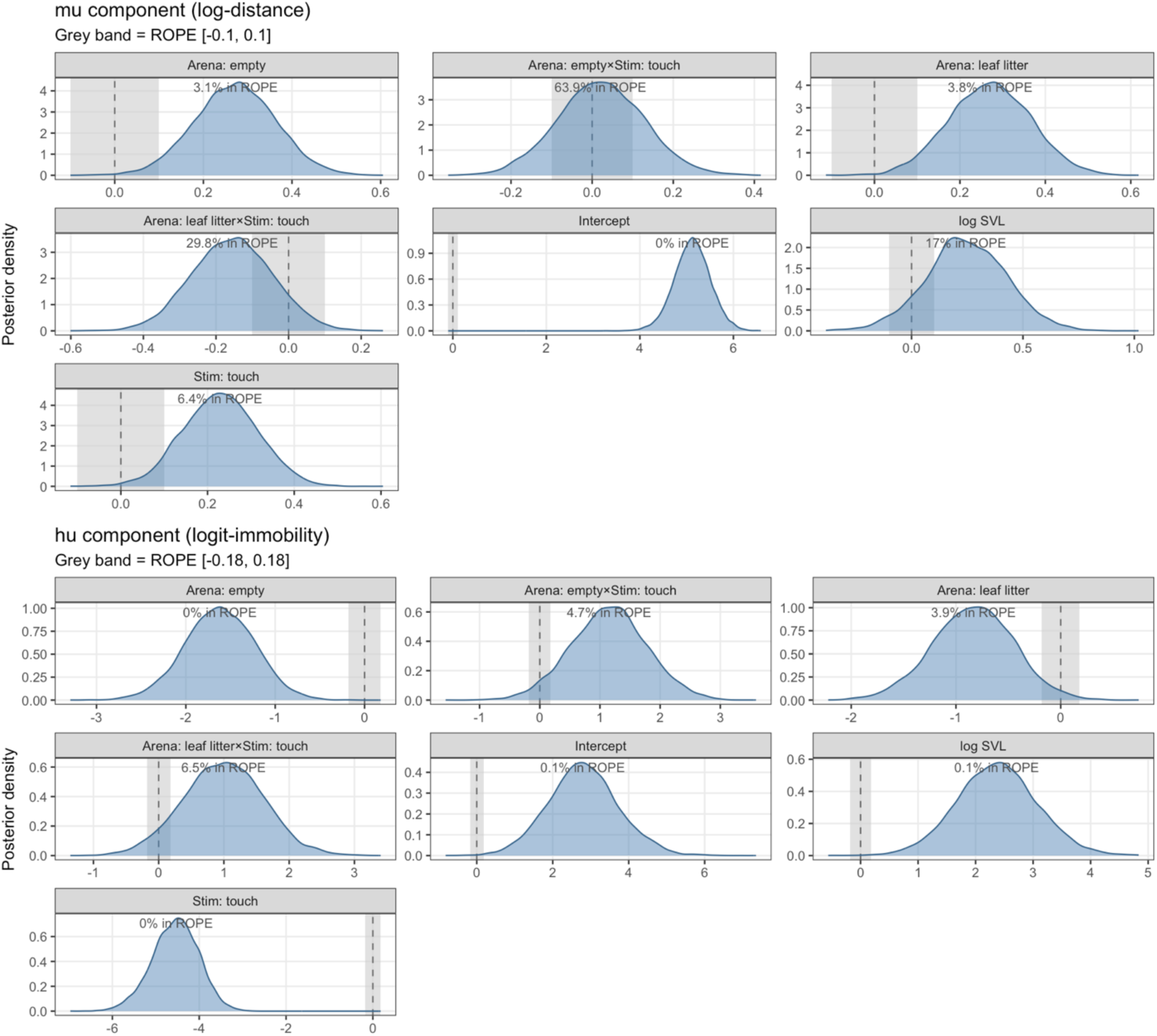
ROPE (Region of Practical Equivalence) and pd (probability of direction) analysis for all fixed effects in both model components. Multi-panel figure showing the posterior distribution of each parameter with the ROPE region highlighted. Parameters with pd > 97% and ROPE < 2.5% are considered to have a clear directional effect.

**Figure S10.**
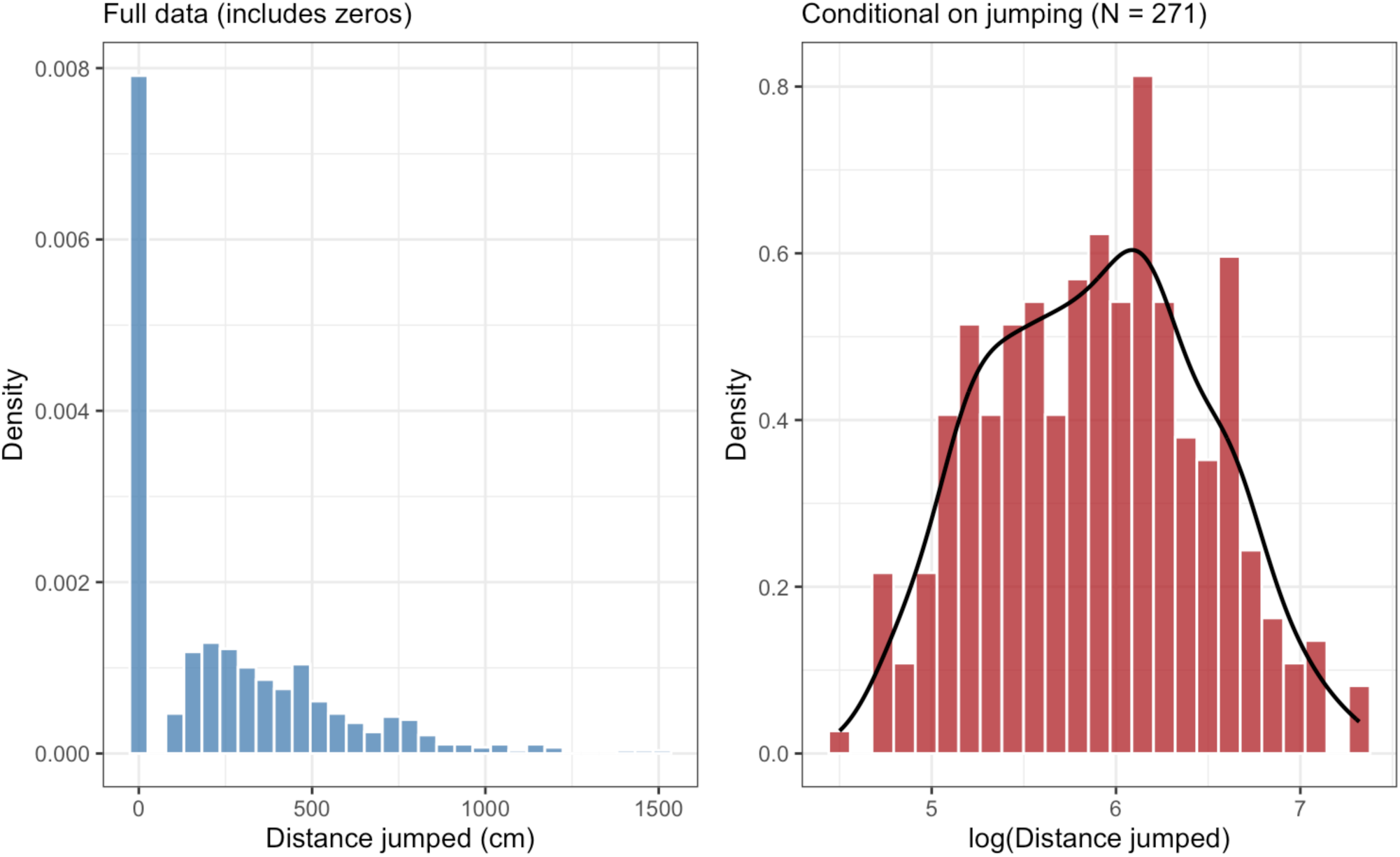
Distribution of the response variable (jump distance) across all 534 trials, showing the bimodal structure with a spike at zero (immobility) and a right-skewed continuous distribution for non-zero values. This structure motivated the hurdle model specification.

**Figure S11.**
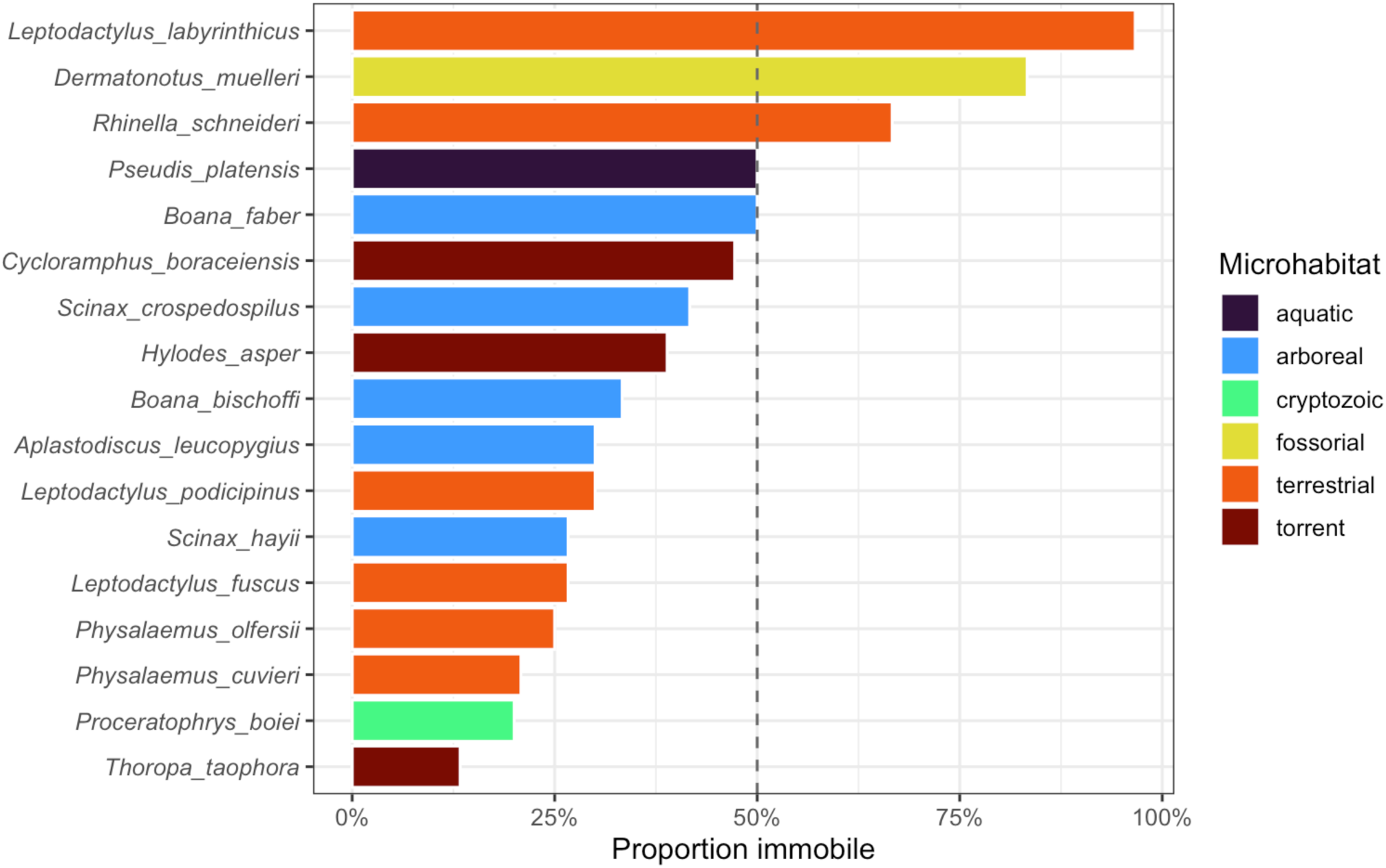
Proportion of zero observations (immobility) by species, ordered by decreasing proportion. Species are colored by microhabitat type. Fossorial and terrestrial species show the highest zero proportions, while arboreal and torrent species show the lowest.

**Figure S12.**
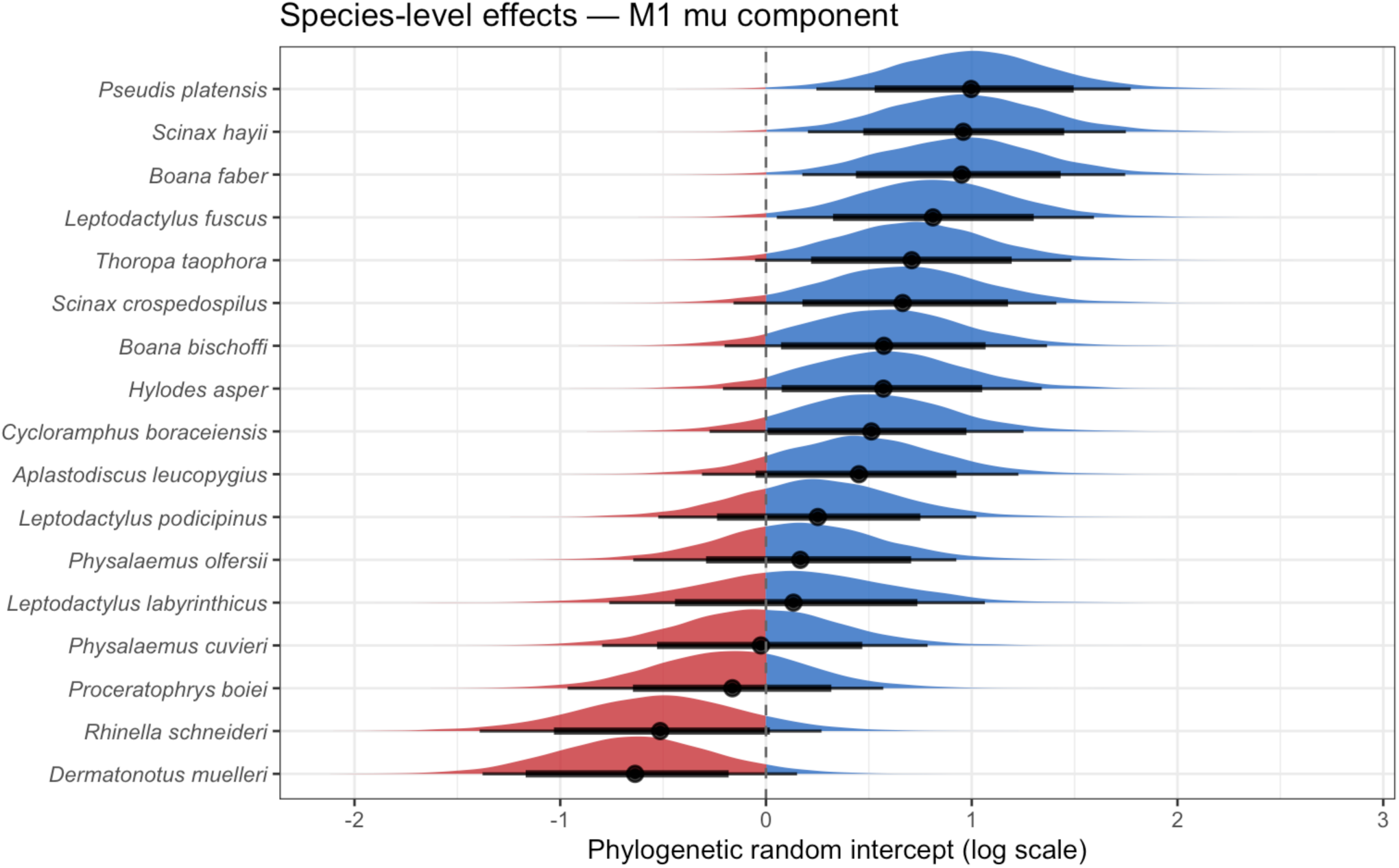
Species-level random effects for both model components. Points and horizontal bars show the posterior median and 95% ETI for each species. Species with ETIs entirely above or below zero differ substantially from the population mean.

**Figure S13.**
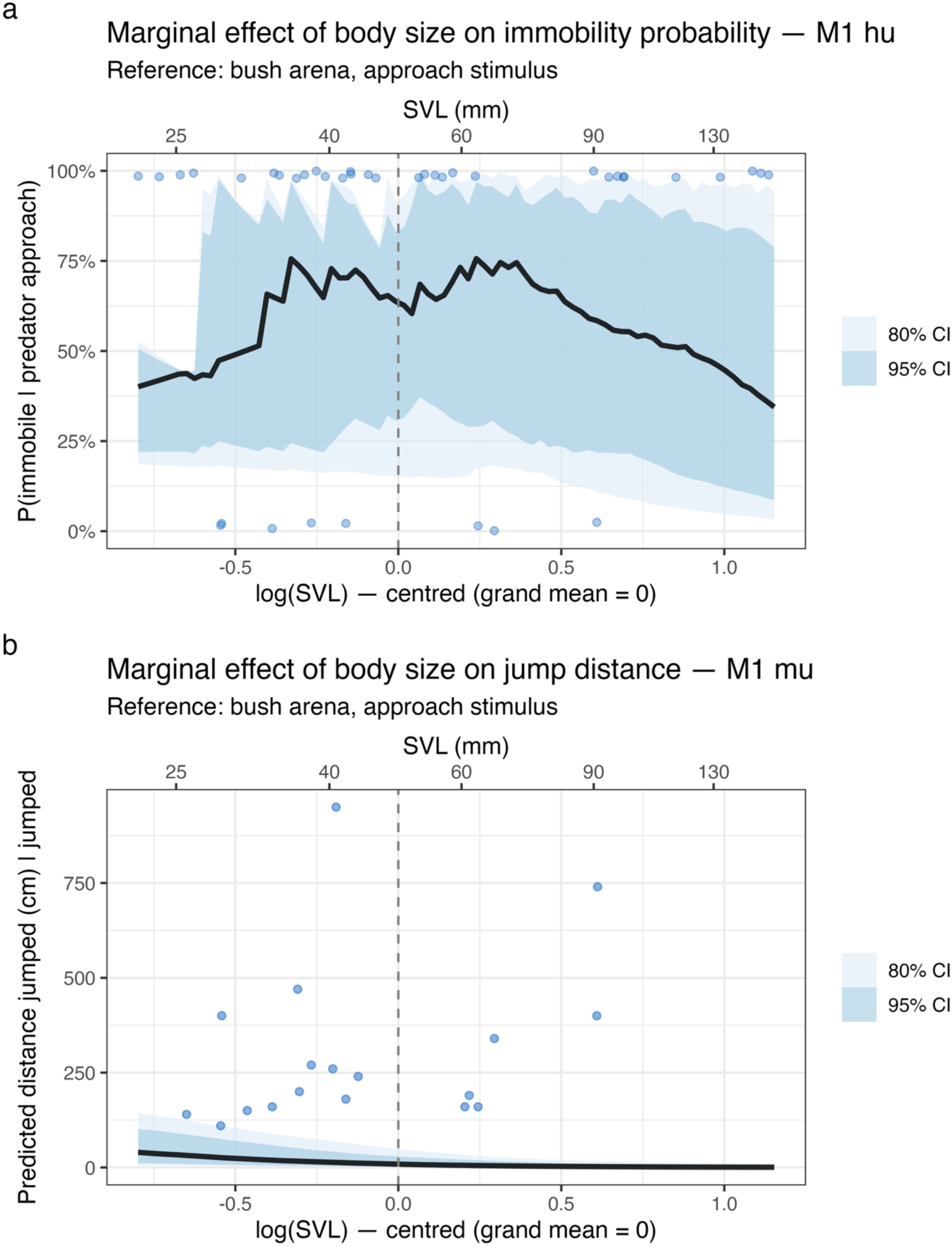
Marginal effect of snout–vent length (SVL) on the two response components. (a) Effect on immobility probability (*hu* component): clear positive relationship. (b) Effect on conditional jump distance (*mu* component): weak positive trend with wide credible interval.

**Figure S14.**
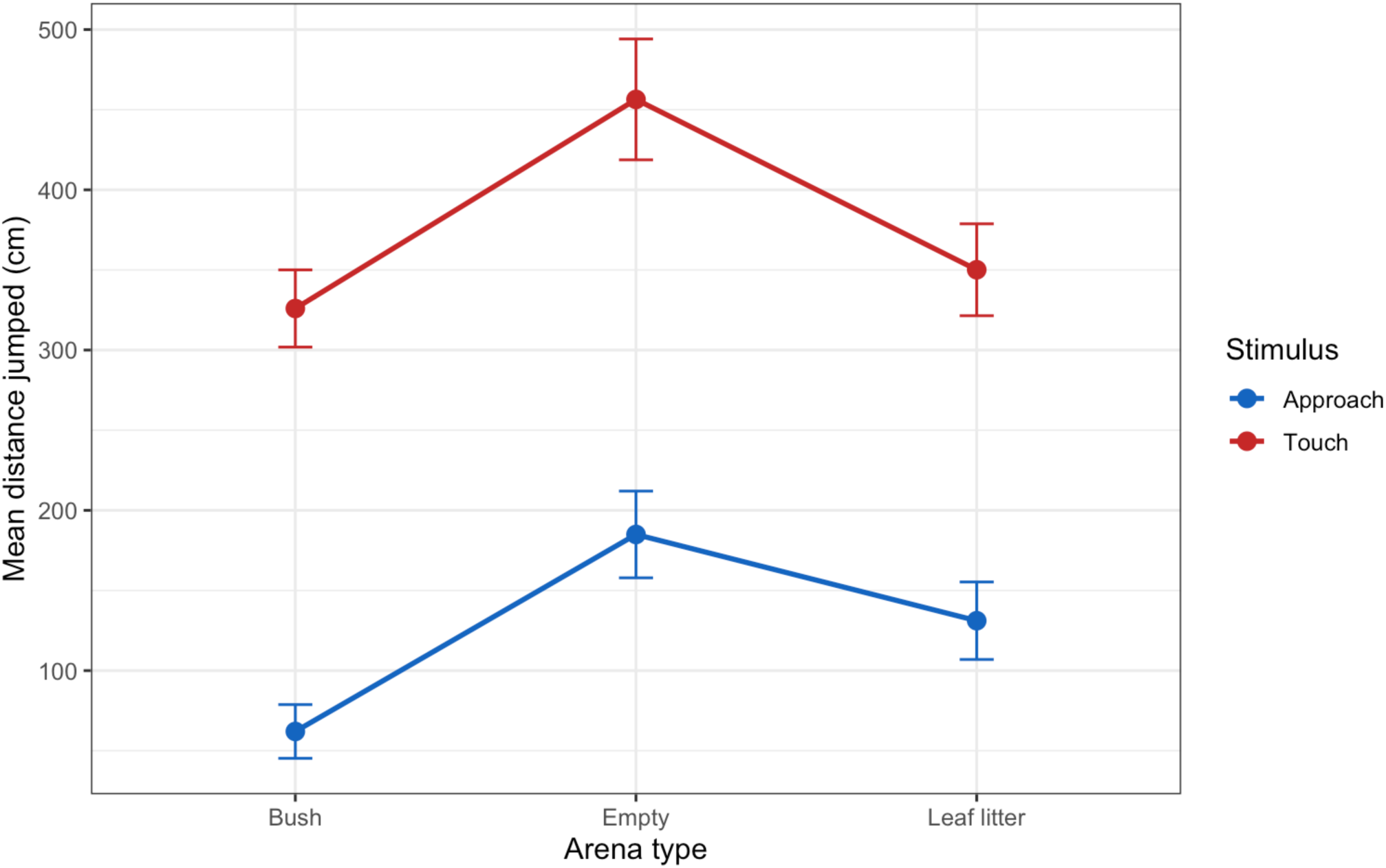
Arena × stimulus interaction effects on the linear predictor scale.

